# FASN Inhibition Resensitizes Chordoma to Radiotherapy by Targeting Adaptive Unsaturated Fatty Acid Metabolism

**DOI:** 10.64898/2026.05.11.724415

**Authors:** Ruolun Wei, Yifan Meng, Emon Nasajpour, Dena Panovska, Helena C.M. Oft, Yao Lulu Xing, Christine K Lee, Juan Carlos Fernandez-Miranda, Matei A. Banu, Richard N. Zare, Claudia K. Petritsch

## Abstract

**SUMMARY:** Chordoma, a rare malignant notochordal tumor of the skull base and spine, is typically resistant to chemotherapy and radiotherapy and exhibits aggressive local recurrence. Here we show that chordoma recurrence correlates with a coordinated upregulation of monounsaturated fatty acids (MUFAs) and polyunsaturated fatty acids (PUFAs), a low PFA/MUFA ratio and an adaptive, lipid peroxidation-resistant state that protects against DNA damage and cell death. Single-cell metabolic profiling identified a tumor subpopulation marked by a fatty acid biosynthesis-high state coupled to stemness. RT-tolerance was directly linked to elevated FASN and lipid droplet (LD) expansion, and MUFA-loading phenocopied RT-tolerance in chordoma cells. Mechanistically, LDs accumulated in response to RT via generation of ROS, and subsequent activation of ER-stress, SREBP1 and Fatty Acid Synthetase (FASN). DESI-MS showed that low-dose irradiation was sufficient to increase MUFAs early and build peroxidation resistant MUFA-LDs, whereas PUFA induction required a higher radiation dose. In a spatially defined manner in a patient-derived xenograft. Finally, in silico knockout and pharmacologic FASN blockade restored radiosensitivity and apoptosis in vitro and in vivo. Collectively, our result support a unifying model in which RT resistance in chordoma is shaped by an adaptive fatty acid metabolic program that buffers oxidative injury and increases survival of RT-resistant, stem-like tumor subpopulations. These findings further support FASN inhibition as a practical radiosensitization strategy for chordoma particulary where RT dose escalation is constrained by anatomy.

**KEYPOINTS:**

1. Recurrent chordoma exhibits fatty acid–associated metabolic reprogramming.
2. MUFA-associated lipid droplet accumulation is linked to radioresistance in chordoma cells.
3. Targeting FASN restores radiotherapy sensitivity of chordoma in vitro and in vivo.

**IMPORTANCE OF STUDY:** This study underscores the clinical importance of targeting metabolic vulnerabilities to restore radiosensitivity in chordoma. By integrating transcriptomics, metabolomics, and in vitro and in vivo models, we identified adaptive fatty acid metabolic reprogramming as a central mechanism of RT resistance in chordoma. Recurrent tumors were characterized by coordinated enrichment of unsaturated fatty acids, especially monounsaturated fatty acids (MUFAs), together with a low PUFA/MUFA ratio and a lipid peroxidation-resistant state. Mechanistically, RT-tolerance chordoma cells exhibited a high-FASN state driven by activation of the ROS–ER stress–PERK/SREBP1/FASN axis, leading to intracellular lipid droplet expansion. Importantly, genetic and pharmacologic inhibition of FASN restored radiosensitivity and enhanced apoptosis in both in vitro and in vivo models, suggesting a translatable therapeutic strategy. Together, these findings link adaptive metabolic reprogramming to RT resistance and support new therapeutic approaches for chordoma management.

## INTRODUCTION

Chordomas are prototypical ultra-rare mesenchymal cancers with an overall incidence estimated at 0.8 cases per million persons annually^1–3^. Their low-grade histopathology is contrasted by aggressive local invasion with frequent early recurrence and late metastases^4^. Originating from remnants of the embryonic notochord within the axial skeleton, chordomas present at nearly equal rates in the skull base, mobile spine, and sacrum^5,6^. Molecularly, chordomas are defined by expression of T-box transcription factor, encoding brachyury (TBXT), which is responsible for notochord development^7–9^.

Standard of care for skull base chordomas (SBC) relies on maximal safe resection followed by adjuvant radiation therapy (RT), while current chemotherapy regimens offer minimal benefit and is not frequently employed^4,10,11^. Moreover, gross total resection is rarely achievable, even with aggressive surgery, given the tumor’s close proximity to critical neurovascular structures^12^. High dose RT would be necessary to suppress SBC recurrence, but its significant morbidity is prohibitive. Not surprisingly loco-regional recurrence rates for SBCs exceed 50%^13^, raising the urgent need for RT-sensitizing strategies. The mechanisms driving RT resistance in chordoma are largely unknown and, thus, drugs that increase RT responses are not used.

Radiation induces DNA damage directly through ionization and indirectly through upregulation of reactive oxygen species (ROS), ultimately leading to cell death. ROS promote lipid peroxidation in response to RT, while simultaneously inducing DNA damage and repair responses. A key determinant of vulnerability to ROS is the ratio of polyunsaturated to monounsaturated fatty acids (PUFA to MUFA ratio): PUFA-containing phospholipids are highly prone to peroxidation and can drive membrane damage and ferroptosis. MUFAs, on the other hand, are relatively peroxidation-resistantand can bufferlipid peroxidation in part by promoting biogenesis of lipid droplets (LD), which are organelles known to reduce oxidative stress^14–17^. Emerging evidence further suggests that RT-induced oxidative stress can activate adaptive anti-apoptotic and anti-ferroptotic programs that support tumor cell survival and RT-resistance in cancer. However, the lipid metabolic programs that shape vulnerability to RT—induced cell death and their contribution to RT resistance remain poorly defined.

Here we identified fatty acid rewiring as an adaptative mechanism to RT, conferring RT tolerance in chordoma. Integrating transcriptomics of patient-derived chordoma, global and spatial metabolomics, and functional models, we show that recurrent chordoma exhibit upregulated fatty acid synthesis, and enrichment of MUFA and, albeit at lower levels, PUFA, resulting in a decreased PUFA/MUFA ratio. Single cell analysis shows elevated fatty acid biosynthesis defines an adaptive stem-like state in tumor cells revealed by metabolic profiling. The ROS-PERK-SREBP1-FASN axis drives adaptive lipid remodeling and enlarged lipid droplets (LDs), conferring RT-tolerance in chordoma model. FASN inhibition suppresses radiation- and MUFA-driven lipid remodeling, restores radiosensitivity *in vitro*, and potentiates the anti-survival effects of low-dose fractionated RT in subcutaneous and orthotopic models. Spatial metabolomics reveals intratumoral fatty acid heterogeneity, with mutually exclusive MUFA and PUFA distributions, and a reduced PUFA/MUFA ratio in the tumor core, consistent with a metabolically buffered niche linked to RT resistance. These findings identify stress-adaptive lipid remodeling as a driver of chordoma RT resistance and nominate FASN inhibition as a practical RT sensitization strategy.

## RESULTS

### Recurrent chordoma exhibits upregulated fatty acid metabolic reprogramming and MUFA enrichment

The mechanisms for RT resistance are associated with high rates of recurrence and are poorly understood. To define features of recurrence, we analyzed 13 clinical specimens from SBC patients by whole transcriptome profiling (Supplementary Table S1A-C). Compared with primary SBCs, recurrent SBCs exhibited increased expression of programs associated with fatty acid metabolism and degradation (Fig. 1A; Supplementary Fig.1) and fatty acid synthase activity (Fig. 1B). Amongst the differentially expressed genes (DEGs) associated with lipid metabolism, transcripts encoding proteins with functions in fatty acid production and metabolic regulation, including *FASN* and *CPT1A* and *1C, SCD, ALOX15, FATP3, NR1H3, CYP11A1*, were consistently elevated in recurrent chordoma (Fig. 1C). Protein–protein interaction (PPI) analysis further placed FASN at a hub position within this metabolic network (Fig. 1D).

**Figure 1.**
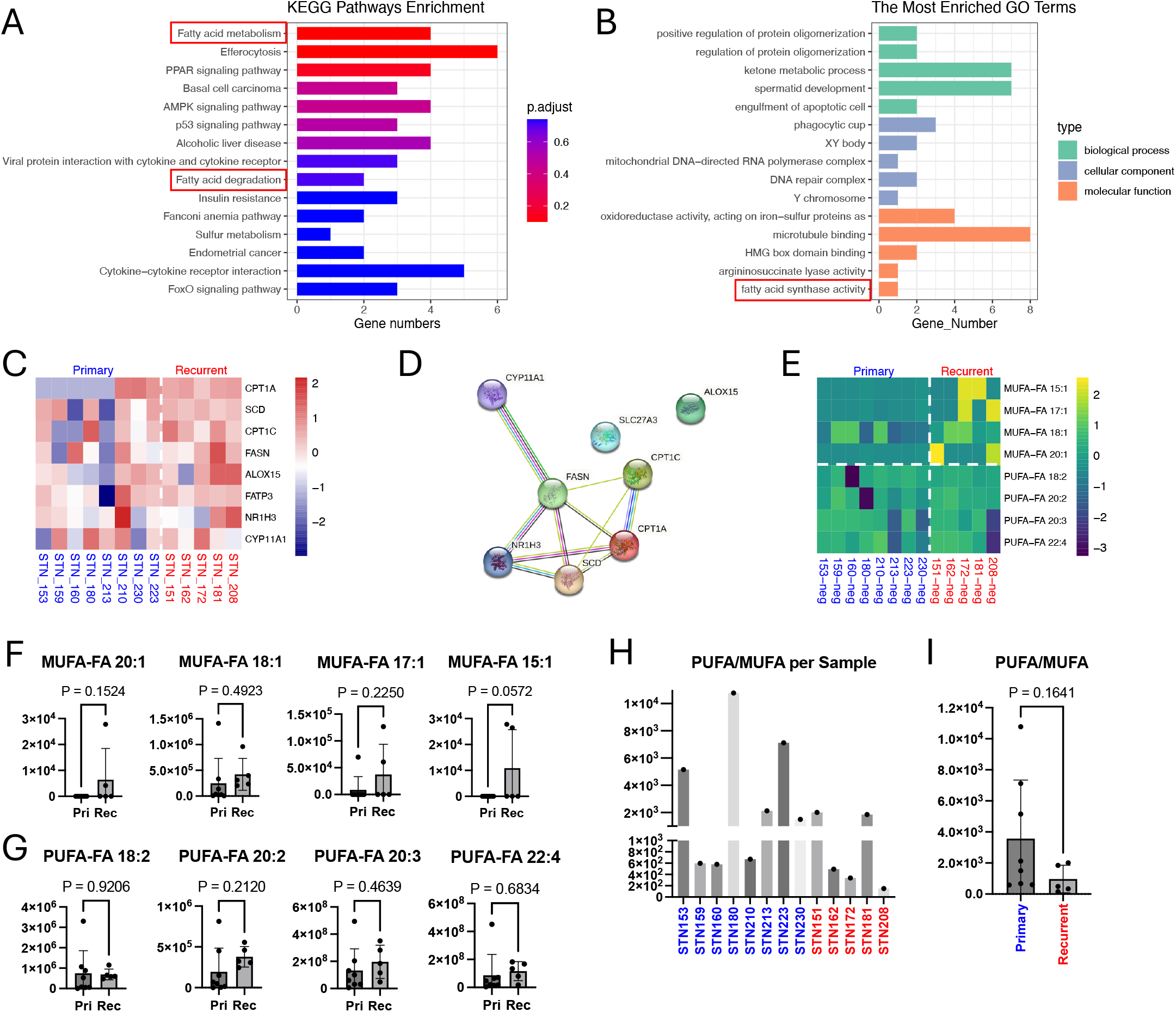
Recurrent chordoma exhibits upregulated fatty acid metabolic reprogramming and MUFA enrichment. (A) Unsupervised KEGG enrichment analysis of DEGs upregulated in recurrent tumors revealed high enrichment of fatty acid metabolism and fatty acid degradation pathways. (B) Unsupervised GO enrichment analysis of DEGs upregulated in recurrent tumors revealed high enrichment of fatty acid synthase activity. (C) Heatmap shows increased expression of fatty acid metabolism–related genes in recurrent tumors. (D) A protein–protein interaction (PPI) network constructed from metabolism-related genes among the upregulated DEGs revealed FASN as a central regulatory hub within the metabolic network. (E) Heatmap of the Nano-ESI-MS analysis revealing a preferential increase in MUFA levels in recurrent tumors, whereas PUFA levels were less frequently elevated. (F) Recurrent tumors exhibit a preferential increase in MUFA levels. Y-axis shows normalized intensity values for the indicated lipid species measured by Nano-ESI MS. (G) PUFA abundance was less increased in recurrent tumors, albeit levels varied amongst tumor samples. Y-axis shows normalized intensity values for the indicated lipid species measured by Nano-ESI MS. (H) Recurrent tumors exhibit a trend toward a reduced PUFA/MUFA ratio compared with primary tumors albeit ratios varied amongst tumor samples. Y-axis shows normalized intensity values for the indicated lipid species measured by Nano-ESI MS. (I) Recurrent tumors overall exhibit a trend towards lower PUFA/MUFA ratio than primary tumors. Y-axis shows normalized intensity values for the indicated lipid species measured by Nano-ESI MS.

Because fatty acid synthesis, particularly driven by FASN activity, contributes to the production of monounsaturated fatty acids (MUFAs)^18,19^, we next performed untargeted nano-ESI mass spectrometry on human SBC specimens with a focus on measuring fatty acid levels. A marked trend towards enrichment of MUFAs was detected in recurrent versus primary SBCs (Figs. 1E, 1F), whereas polyunsaturated fatty acids (PUFAs) also upregulated and showed less pronounced and less frequent differences between primary and recurrent SBC (Figs. 1E, 1G).

We next examined the ratio of PUFA to MUFA as a key determinant of oxidative vulnerability^20,21^. Primary SBCs displayed a relatively higher PUFA/MUFA ratio than recurrent SBCs (Figs. 1H, 1I).

### Elevated fatty acid biosynthesis defines an adaptive stem-like state in tumor cells revealed by single cell metabolic profiling

To characterize fatty acid metabolism further and assess how it affects intra-tumoralheterogeneity, we analyzed a public dataset of single-cell transcriptomic data from 4 SBC samples (Fig. 2A). *TBXT+* tumor cells were extracted based on established chordoma markers and re-clustered for higher-resolution analysis (Fig. 2B). This analysis resolved four distinct tumor cell states (sc_0–sc_3). Metabolic pathway profiling using scMetabolism revealed a prominent fatty acid biosynthesis program in sc_1, whereas sc_0 and sc_2 showed comparable enrichment of glycolysis and oxidative phosphorylation signatures. In contrast, sc_3 was characterized by a strong cytochrome P450–associated metabolic program (Fig. 2C). Notably, the enrichment of fatty acid biosynthesis in sc_1 is consistent with activation of fatty acid biosynthesis pathways, including FASN-driven lipid synthesis, suggesting that this cell state promotes adaptive fatty acid remodeling in recurrent tumor.

**Figure 2.**
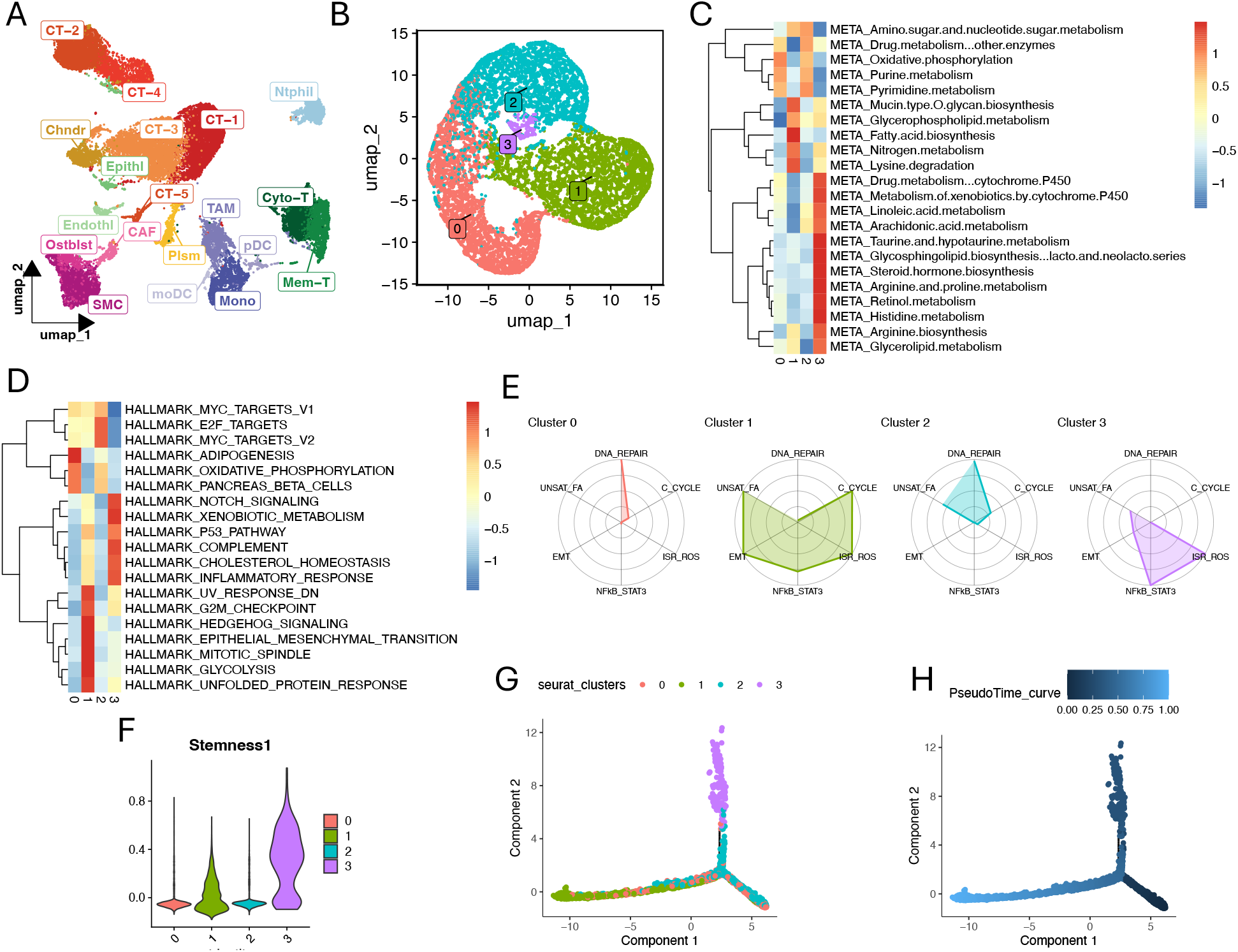
Elevated fatty acid synthesis defines an adaptive stem-like state in tumor cells revealed by metabolic profiling. (A) Quality control, clustering, and cell-type annotation of SBC single-cell RNA-seq dataset. (B) *TBXT+* tumor compartment extraction based on established chordoma markers, followed by re-clustering, which revealed four subclusters (sc_0–sc_3; resolution = 0.05). (C) scMetabolism-based pathway activity profiling across tumor subclusters. (D) Single-cell HALLMARK pathway enrichment across tumor subclusters. (E) Radar plot summarizing curated stress- and resistance-relevant gene modules (DNA repair, cell cycle [c_cycle], ROS/stress response [ISR_ROS], STAT3 signaling, EMT, and unsaturated fatty acid metabolism [UNSAT_FA]) across tumor subclusters. (F) Stemness1 module score based on canonical stem-like genes (PROM1, SOX2, NANOG, KLF4, MYC, ALDH1A1, NES, ITGA6) across tumor subclusters. (G) Monocle pseudotime trajectory inference of tumor subclusters. (H) Monocle pseudotime distribution indicating sc_1 is positioned toward the late stage of the inferred trajectory.

To further define functional differences across cell states, we performed single-cell hallmark pathway enrichment analysis. This analysis revealed that the sc_1 subcluster was enriched for proliferative programs, including Hedgehog signaling, epithelial–mesenchymal transition (EMT), and mitotic spindle pathways, together with increased UV response–down features (Fig. 2D). These findings link fatty acid biosynthesis activity with proliferative and invasive tumor cell states, suggesting that fatty acid remodeling may support tumor growth and promote tumor progression.

To summarize key stress-response and resistance-relevant programs, we curated gene signatures spanning DNA repair (DNA_REPAIR), cell cycle (C_CYCLE), ROS/stress response (ISR_ROS), STAT3 signaling (NFkb_STAT3), EMT (EMT), and unsaturated fatty acid metabolism (UNSAT_FA), and quantified these modules using radar plot. In addition to elevated unsaturated fatty acid biosynthesis, sc_1 scored highly for EMT, cell-cycle activity, and ROS/stress response (Fig. 2E). Since stemness is associated with cell plasticity and therapy resistance, we then constructed a Stemness1 module using canonical tumor stem-like genes (“PROM1”, “SOX2”, “NANOG”, “KLF4”, “MYC”, “ALDH1A1”, “NES”, “ITGA6”). While sc_3 appeared most stem-like based on this score, sc_1 also showed increased stem-like features relative to other subclusters (Fig. 2F). Monocle-based pseudotime inference positioned sc_1 late in the trajectory of tumor cell evolution (Figs. 2G, 2H), suggesting that this subpopulation characterized by elevated fatty acid biosynthesis represents an adaptive state acquired by more mature tumor cells, rather than a therapy-independent intrinsic stem-like state.

Together, these results indicate that the fatty acid biosynthesis–high sc_1 state represents a more mature, stress-adaptive tumor cell state, with concurrent proliferative and stemness features.

### PERK-SREBP1-FASN axis drives adaptive lipid remodeling conferring RT-tolerance in chordoma

To investigate whether elevated fatty acid biosynthesis constitutes an adaptation to RT, we repeatedly irradiated a patient-derived SBC cell line CH22 to generate an RT-resistant chordoma cell line (CH22R). Additionally, we modeled a MUFA-high phenotype by supplementing CH22 cells with oleic acid (CH22OA, Fig. 3A). CH22R cells exhibited elevated Fatty Acid Synthetase (FASN) expression, even after passaging *in vitro*, establishing elevated fatty acid biosynthesis as an adaptive response to RT (Fig. 3B). Given the role of the FASN–MUFA axis LD biogenesis^14–17^, we next assessed changes in intra-cellular lipid storage by BODIPY staining and confocal microscopy. CH22R and CH22OA both displayed significantly enlarged LDs compared with parental CH22 cells (Fig. 3C). Consistent with acquired RT tolerance, a single dose of irradiation (6 Gy) induced significantly lower γH2AX foci, as a conduit for DNA damage, in CH22R and CH22OA versus parental CH22 cells (Fig. 3D). Moreover, CH22OA showed reduced GADD45A expression, and diminished cleavage of PARP1 and caspase-3 relative to parental CH22 cells, supporting the interpretation that LD accumulation attenuates DNA damage responses and decreases cell death following RT (Fig. 3E).

**Figure 3.**
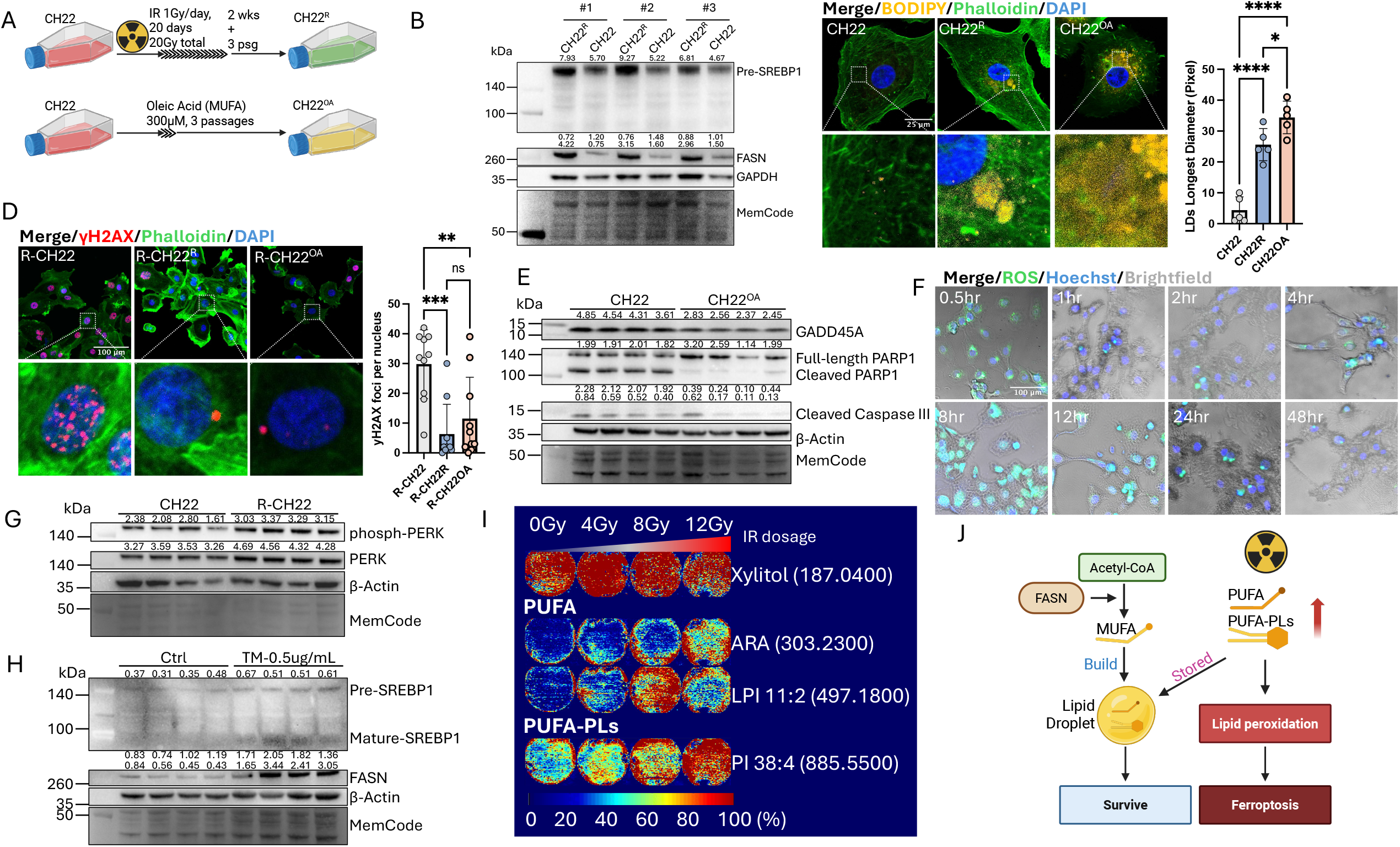
The PERK-SREBP1-FASN axis drives adaptive lipid remodeling conferring RT-tolerance in chordoma. (A) Schematic of two complementary SBC models of RT adaptation and mono-unsaturated fatty acids (MUFA): a radiation-adapted line generated by repeated *in vitro* irradiation (CH22^R^), and an oleic acid– treated line modeling MUFA-driven RT responses (CH22^OA^). (B) Immunoblot showing increased FASN and pre-SREBP1 protein expression in CH22^R^ cells, relative to the parental CH22 cells. Quantitative data of each band were acquired using Image Lab software (Bio-Rad). (C) BODIPY staining and confocal images showing significantly enlarged lipid droplets (LDs) in CH22^R^ and CH22^OA^ compared with parental CH22 cells. (62× oil-immersion objective, nuclei stained with DAPI). (D) Immunocytochemistry showing significantly lower γH2AX (phospho-Ser139) foci in CH22^R^ and CH22^OA^ after single-dose IR (6 Gy) compared to parental CH22 cells. (20× objective, nuclei stained with DAPI). (E) Immunoblot showing lower induction of GADD45A and reduced cleavage of PARP1 and caspase-3 in CH22^OA^, compared with parental CH22, following a single dose of irradiation (6 Gy). (F) Live-cell tracking of intracellular ROS dynamics in CH22 after a single dose of irradiation (6 Gy) showing an early peak at 0.5h, a transient dip at 1–4h, a second peak at 8–12 hr, and a decline by 24–48 hr. (20× objective, nuclei stained with DAPI). (G) Immunoblot analysis of ER-stress signaling in untreated (CH22) and irradiation-treated (R-CH22) CH22 cells, showing early PERK phosphorylation detectable at 6 h post-irradiation (6 Gy), accompanied by increased total PERK protein levels. (H) immunoblot analysis of CH22 cells treated with tunicamycin (TM), showing that TM-induced ER stress elevates levels of precursor SREBP1 (pre-SREBP1) and mature (mSREBP1) SREPB1 and FASN as a downstream effector of mSREBP1. (J) DESI–MS of adherent live CH22 cell (grown on coverslips), using xylitol as a stable internal reference for cell distribution, revealed a dose-dependent accumulation of in PUFA and PUFA-PL species in 12 h after irradiation, consistent with irradiation-associated expansion of the cellular PUFA pool. (color bar (0– 100%) reports relative abundance with warmer colors corresponding to higher concentrations). (J) Schematic of model: RT-induced ROS and ER stress activate the PERK–SREBP1–FASN axis, driving MUFA-enriched neutral lipid biosynthesis and LD accumulation. This adaptive lipid remodeling sequesters excess PUFAs and PUFA-PLs, thereby limiting lipid peroxidation and cell death and conferring RT tolerance.

To identify upstream drivers of this adaptive fatty acid biosynthesis reprogramming, we examined RT-induced early stress responses. Because ionizing radiation is known to rapidly generate ROS^22^, we monitored ROS dynamics in CH22 cells. Interestingly, we observed a biphasic pattern: an early peak at 0.5 h, a reduction at 1–2 h, and a second peak at 8–12 h post-irradiation (Fig. 3F). ROS can trigger endoplasmic reticulum (ER) stress and activate the unfolded protein response, including phosphorylation of Protein kinase RNA-like ER kinase (PERK). Accordingly, PERK phosphorylation was detectable as early as 6 hours after irradiation (6 Gy), accompanied by an increase also in total PERK protein levels, in CH22 cells (Fig. 3G). PERK is known to activate sterol regulatory element-binding protein 1 (SREBP1), a transcription factor that drives expression of *FASN* and other lipogenic genes under lipid insufficiency or ER stress ^23,24^. Irradiation induced higher levels of the inactive form of SREBP1 (pre-SREBP1) in CH22R cells compared with CH22 parental cells (Fig. 3B). Moreover, because FASN-driven *de novo* fatty acid biosynthesis has been linked to RT resistance across multiple cancers, we asked whether PERK activation induces the SREBP1/FASN axis in CH22 cells^25–27^. Indeed, treatment with the PERK activator tunicamycin increased SREBP1 protein levels and maturation of pre-SREBP1, and elevated FASN expression, as a known downstream effector of mature SREBP1 (mSREBP1) (Fig. 3H).

RT can expand the cellular PUFA pool, promoting lipid peroxidation, a process where ROS oxidize PUFAs in the cell and at organelle membranes, damaging cell membranes and causing cell death by apoptosis and ferroptosis^28^. We therefore profiled PUFA and PUFA phospholipids (PUFA-PLs) species in CH22 cells by DESI-MS at 12h post-irradiation. Both PUFA and PUFA-PL species increased in a dose-dependent manner, indicating irradiation-driven early expansion of a lipid pool that is sensitive to lipid peroxidation and cell death (Fig. 3J). Together, these data support a model in which RT-induced ROS and ER stress activate the PERK-SREBP1-FASN axis, driving MUFA-enriched neutral lipid biosynthesis and LD accumulation. Adaptive increases in MUFA, as detected by enlarged LDs, sequester excess PUFAs and PUFA-PLs, thereby limiting lipid peroxidation-induced cell death and conferring RT tolerance (Fig. 3I).

### Spatial MUFA/PUFA distributions reveals intratumoral heterogeneity of redox-buffered niches

Next, we assessed the intratumoral heterogeneity of chordomas to define potential spatial aspects of lipid peroxidation. We determined the spatial distribution of MUFAs and PUFAs in a treatment-naïve patient-derived xenograft (PDX) mouse model of SBC using DESI-MS. Bright-field imaging guided sectioning through the largest tumor plane, encompassing the injection tract (Fig. 4A). Tumor boundaries were defined on H&E-stained sections (Fig. 4B); Ki-67 revealed a proliferative rim at the tumor periphery and cleaved caspase-3 immunostaining identified scattered apoptotic foci within the core (Fig. 4C, 4D). Xylitol served as a reference for overall cellular distribution (Fig. 4E), and Hex2Cer, an abundant sphingolipid in normal brain tissue, delineated the tissue edge (Fig. 4F).

**Figure 4.**
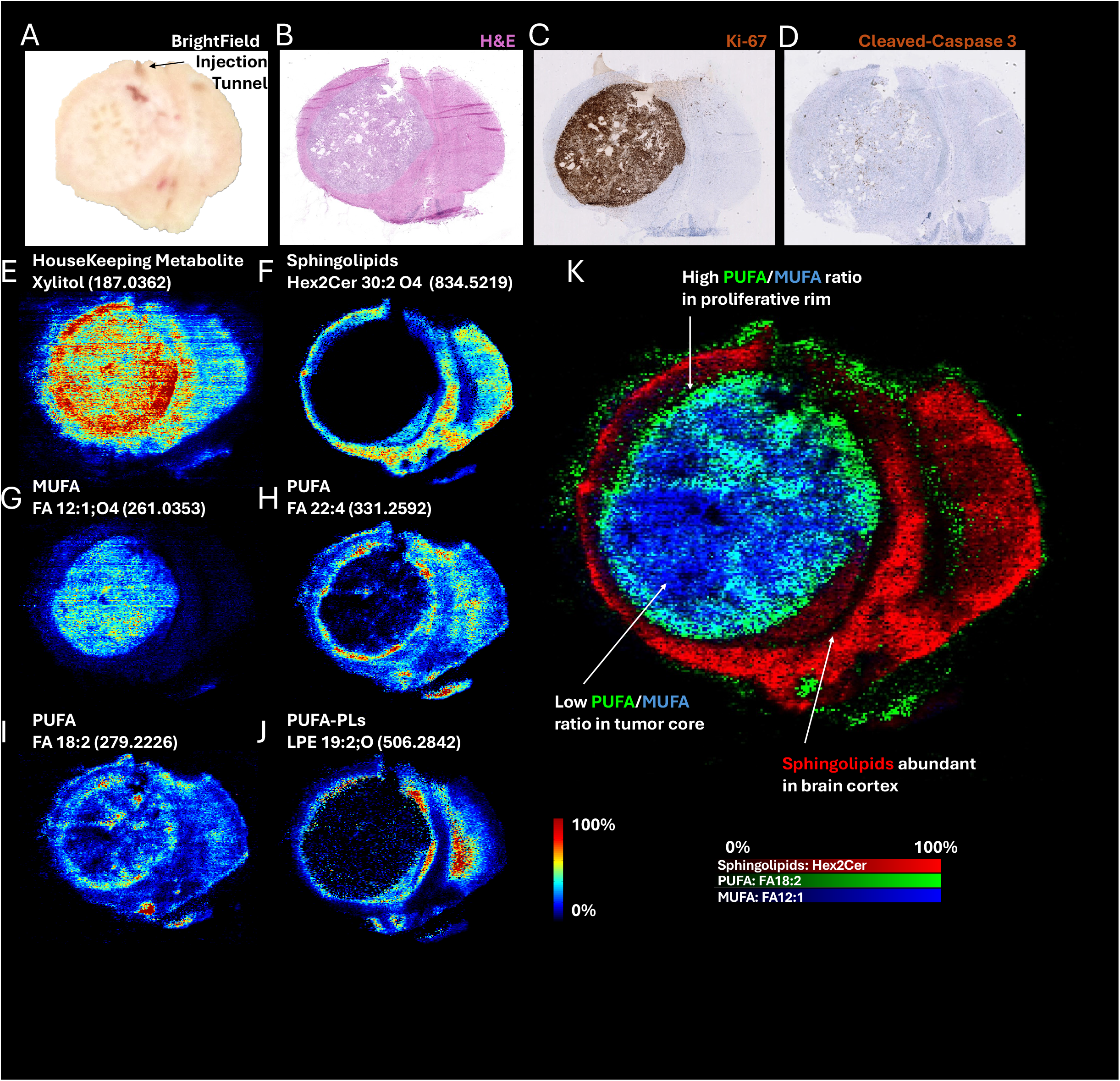
*Spatial MUFA/PUFA distributions reveals intratumoral heterogeneity of redox-buffered niches*. (A-D) Low-magnification brightfield image (A), H&E (B), and matched IHC for Ki-67 (C) and cleaved caspase-3 (D) from a treatment naive mouse bearing an orthotopic CH22 chordoma xenograft. Ki-67 highlights a proliferative rim at the tumor periphery, whereas cleaved caspase-3 marks scattered intratumoral positive foci. Consecutive sections through the tumor center containing the injection tract were used for H&E, IHC, and spatial DESI-MS. (E-K) Xylitol (m/z 187.0362) (E) and Hex2Cer (m/z 834.5219) (F) were used as reference ions in DESI-MS to indicate overall cellular distribution and to delineate the tumor/brain boundary, respectively. DESI-MS spatial metabolomic maps show that MUFAs are enriched in the tumor core and progressively decrease toward the periphery (G) whereas PUFAs (H, I) and PUFA-PLs (J) are enriched in the tumor proliferative rim and adjacent normal brain tissue. Color bar (0–100%) indicates relative abundance; warmer colors = higher signal.

Interestingly, DESI-MS imaging revealed largely spatially segregated MUFA (Fig. 4G) and PUFA (Fig. 4H, 4I, 4J) distributions: regions enriched for MUFAs displayed low PUFA signals, and *vice versa*. The PUFA/MUFA ratio was lowest in the established tumor core, and toward tumor periphery the PUFA/MUFA ratio increased, peaking at the outer proliferative rim. This gradient suggests that MUFA accumulation is a gradual process associated with tumor maturation rather than proliferation. Mechanistically, the low PUFA/MUFA ratio in the core of this untreated chordoma is consistent with a MUFA-skewed lipid state that stabilizes membranes and raises the threshold for lipid peroxidation–mediated cell death. Together, these data indicate that the tumor core is rich in MUFAs that may confer resistance to RT-induced oxidative stress, lipid peroxidation, and cell death.

### In silico and pharmacological FASN targeting reverses RT-driven lipid remodeling and overcomes RT resistance in chordoma cells

Given the central role of FASN-driven MUFA synthesis and RT resistance, we next determined whether it plays a critical role in the adaptive fatty acid synthesis program and could therefore represent a therapeutic vulnerability in SBC. We first predicted the effects of FASN perturbation by applying the *in silico* knockout approach scTenifoldKnk to the single-cell RNAseq data mentioned above. The virtual knockout of FASN in SBC cell subclusters resulted in upregulation of apoptosis-related programs (Fig. 5A), accompanied by downregulation of metabolic pathways and the stemness-associated NOTCH signaling program (Fig. 5B), supporting FASN as a feasible therapeutic vulnerability in SBC.

**Figure 5.**
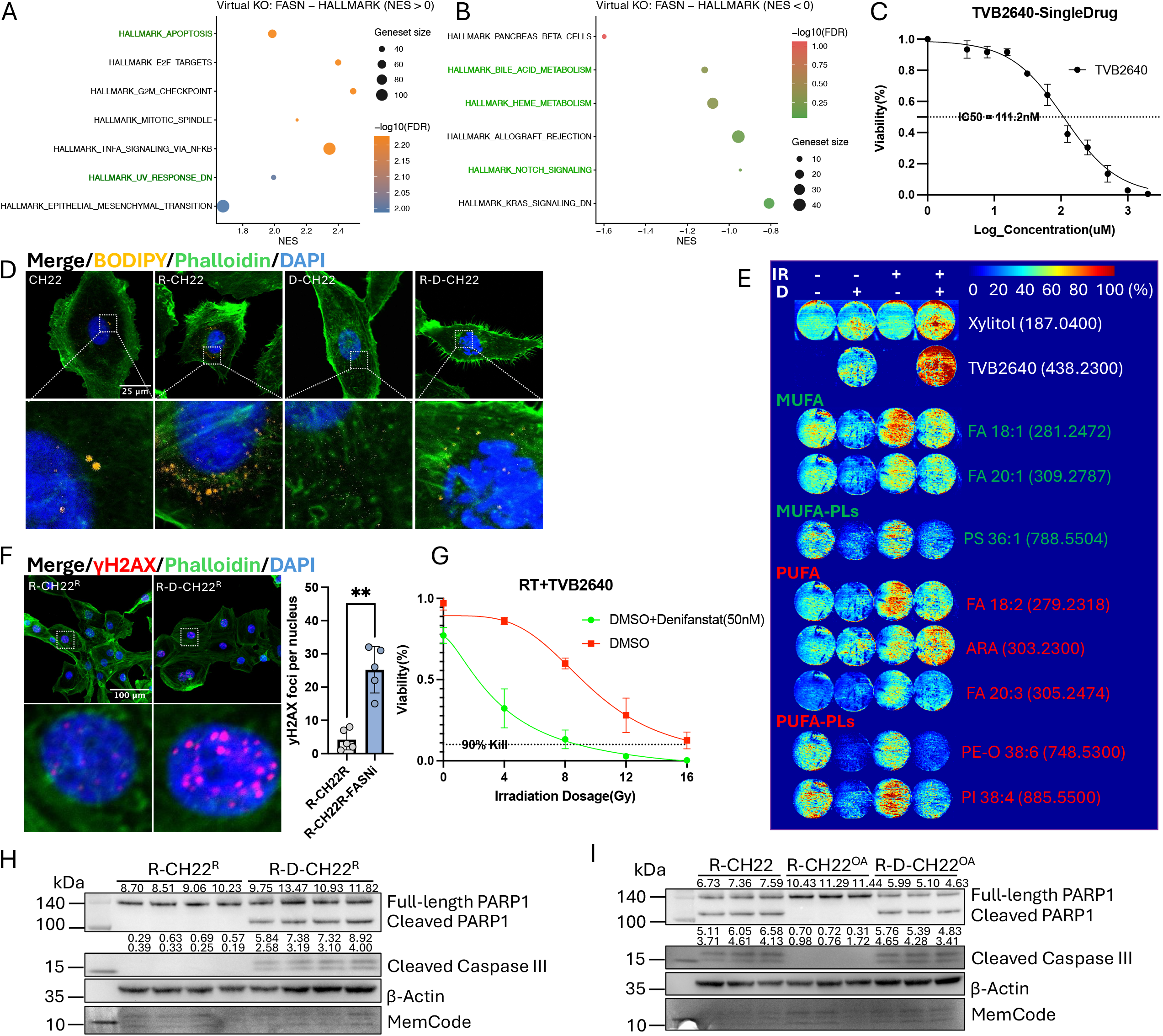
*In silico and pharmacological FASN inhibition reverses RT-driven lipid remodeling and overcomes radiotherapy resistance in chordoma* cells. (A) scTenifoldKnk *in silico* knockout of FASN in chordoma tumor cell clusters showing upregulation of apoptosis-related programs in Hallmark analysis. (B) scTenifoldKnk results showing downregulation of metabolic pathways and the stemness-associated NOTCH signaling program following virtual FASN knockout. (C) Dose–response (IC_50_) curve for TVB-2640 monotherapy in CH22 chordoma cells; nonlinear regression estimates IC_50_ = 111.2 nM. (D) Confocal microscopy showing that TVB-2640 reduces control and irradiation-associated lipid accumulation in CH22 cells. (E) DESI-MS quantification of lipid features following TVB-2640 treatment under baseline and post-irradiation conditions, showing reduced MUFAs and MUFA-containing lipids, minimal change in free PUFA features, and a decreasing trend in irradiation-induced PUFA-containing lipid species with FASN inhibition (color bar (0–100%) reports relative abundance with warmer colors corresponding to higher concentrations). (F) Immunofluorescence and quantification of γH2AX foci showing significantly increased nuclear γH2AX accumulation in CH22^R^ cells after combining irradiation (6 Gy, single dose) with TVB-2640 treatment (20× objective, nuclei stained with DAPI). (H) Immunoblot showing that TVB-2640 resensitized CH22^OA^ cells to low-dose ionizing radiation (1 Gy, single dose), as evidenced by increased cleaved PARP1 and cleaved caspase-3. (I) Immunoblot showing that TVB-2640 resensitized the radioresistant CH22^R^ line to low-dose radiation (1 Gy, single dose), as indicated by induction of cleaved PARP1 and cleaved caspase-3.

Next, we targeted FASN in RT-resistant CH22R cells with TVB-2640, a clinical-stage small molecule inhibitor currently in clinical trial stage. TVB-2640 monotherapy yielded an IC_50_ of approximately 111.2 nM in CH22R cells (Fig. 5C). TVB-2640 also antagonized LD enlargement exacerbated by irradiation (Fig 5D), and effectively reduced MUFAs and MUFA-containing lipids in CH22R cells at baseline and post-irradiation (Fig. 5E). In contrast, PUFA features were not substantially altered by FASN inhibition, as shown by DESI-MS. Interestingly, however, PUFA-containing phospholipid species, particularly those induced after irradiation, showed a decreasing trend with TVB-2640 treatment, paralleling the reduction observed for MUFAs and MUFA-containing lipids (Fig. 5E).

TVB-2640 significantly increased post-irradiation DNA damage in CH22R cells (Fig. 5F). Combined TVB-2640 and low-dose irradiation (4 Gy) eliminated 90% of CH22R cells, an effect equivalent to a 16 Gy single-fraction dose (Fig. 5G). TVB-2640 also reversed oleic acid–induced RT tolerance, with cleaved PARP1 and cleaved caspase-3 detectable as early as 2 h after 2 Gy irradiation (Fig. 5H–I). Together, these data demonstrate that pharmacological FASN inhibition reduces LD accumulation and impairs DNA damage repair capacity, thereby re-sensitizing RT-resistant chordoma cells to radiotherapy.

### FASN inhibition combined with low-dose fractionated RT blocks chordoma growth in subcutaneous model

We next assessed whether the combinatorial pro-apoptotic effects of pharmacologic FASN inhibition with low-dose RT *in vitro* would translate into reduced tumor growth *in vivo*. Subcutaneous CH22 xenograft-bearing mice were randomized into a control, RT alone (RT), TVB-2640 (D) alone, and RT + TVB-2640 (RTD) arm. Mice were treated with low-dose fractionated radiotherapy (LDFR; 1 Gy/day × 4 days; total 4 Gy), TVB-2640 once daily by oral gavage at 25 mg/kg body weight, or a combination of both, starting at 120 mm^3^ tumor volume. Tumor size was measured as described (Methods) (Fig. 6A, Supplemental Fig. S3, Supplemental Fig. S4)^29,30^. LDFR alone did not significantly suppress tumor growth compared to control treatment.TVB-2640 monotherapy slowed tumor growth during treatment but tumors rebounded rapidly after treatment cessation. In contrast, the combination of TVB-2640 and LDFR blocked tumor growth more durably, suppressing growth after treatment cessation (Fig. 6B), as confirmed by resected tumor wet weight and size at endpoint (Figs.6C–6D). TVB-2640 induced on-treatment weight loss ranging from 6%-10%, and weight recovered after treatment cessation, which may indicate an on-target pharmacologic effect (Fig. 6E). Combined RT and FASN inhibition increased cleaved caspase-3 and reduced Ki-67 compared with either monotherapy (Fig. 6F), consistent with both RT sensitization and cytostasis. Collectively, these data demonstrate that FASN blockade converts a subtherapeutic RT regimen into an effective anti-tumor response *in vivo*, albeit with treatment-associated weight loss that should be assessed by tolerability studies in the future.

**Figure 6.**
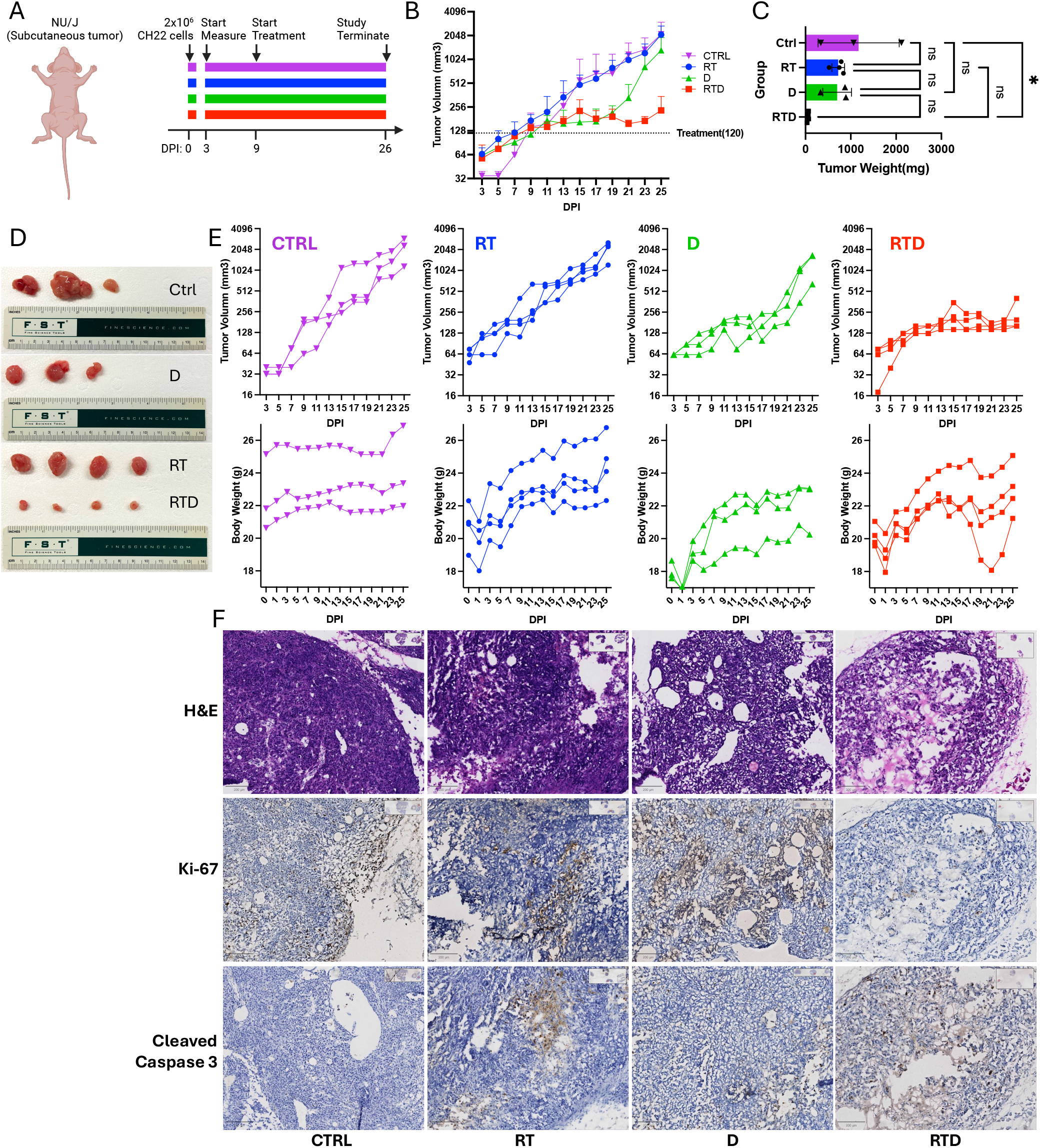
FASN inhibition plus low-dose fractionated RT blocks subcutaneous chordoma growth. (A) Schematic of the subcutaneous chordoma xenograft and treatment timeline with four groups: CTRL (vehicle control); RT (low-dose fractionated radiotherapy, LDFR alone: 1 Gy per day for 4 days); D (TVB-2640 monotherapy, 25 mg/kg per day for 10 days); and RTD (LDFR + TVB-2640 combination). Each group included at least 3 mice (n ≥ 3). (B) Tumor growth curves with the y-axis showing log2-transformed tumor volume (in mm3) and the x-axis showing days post injection (dpi). (C) Endpoint tumor wet weight: the CTRL group was higher than RT and D, and significantly higher than RTD; the RTD combination yielded the lowest tumor weight. (D) Freshly excised tumors of all groups, with a ruler showing inches and centimeters for scale. (E) Tumor growth and body-weight trajectories for CTRL, RT, D, and RTD groups. Tumor curves are plotted with the y-axis as log_2_-transformed tumor volume (mm^3^); body-weight curves use grams (g) on the y-axis; the x-axis shows days post injection (dpi). (F) Representative H&E and matched IHC for Ki-67 and cleaved caspase-3 across treatment groups. In CTRL tumors, Ki-67 staining is high while cleaved caspase-3 is largely absent; in the RTD group (RT + TVB-2640), Ki-67–positive cells are nearly absent, whereas widespread cleaved caspase-3 staining is evident (scale bar as indicated).

### FASN inhibition combined with low-dose fractionated RT blocks chordoma growth in an orthotopic model

Based on our data in the flank chordoma model, we hypothesized that lower-dose fractionated radiotherapy (LDFR) augmented by FASN inhibition could achieve tumor control for SBC xenografts in a more native microenvironment. Orthotopic chordoma were grown near the skull base of immunocompromised mice. Once established, mice were stratified into small-, medium-, and large-size cohorts by 7T MRI on day 20 post-implantation, modeling different tumor stages and taking into account that larger tumor volume was associated with poor outcomes after standard treatment in patients with SBC^1,6,12^. LDFR+TVB-2640 combination treatment (Fig. 7A, Supplemental Figs. S6, Fig. S7). significantly prolonged survival versus vehicle-treated control (log-rank P = 0.0246) (Fig. 7B). Body weight declined transiently during dosing and recovered after treatment cessation, similar to the subcutaneous model (Fig. 7C). Serial MRI before treatment and at end of therapy showed effective growth suppression in mid- and large-sized tumors, characterized by intratumoral cystic/vacuolated changes without expansion of the solid component; in small-size tumors, LDFR+TVB-2640 produced near-complete responses (Fig. 7D). Ki-67 and cleaved caspase-3 staining confirmed that LDFR+TVB-2640 reduced proliferation and induced apoptosis in vivo (Fig. 7E). Together, these data demonstrate that FASN inhibition converts a subtherapeutic RT regimen into effective tumor control in an orthotopic chordoma model, providing preclinical rationale for combining TVB-2640 with radiotherapy in patients.

**Figure 7.**
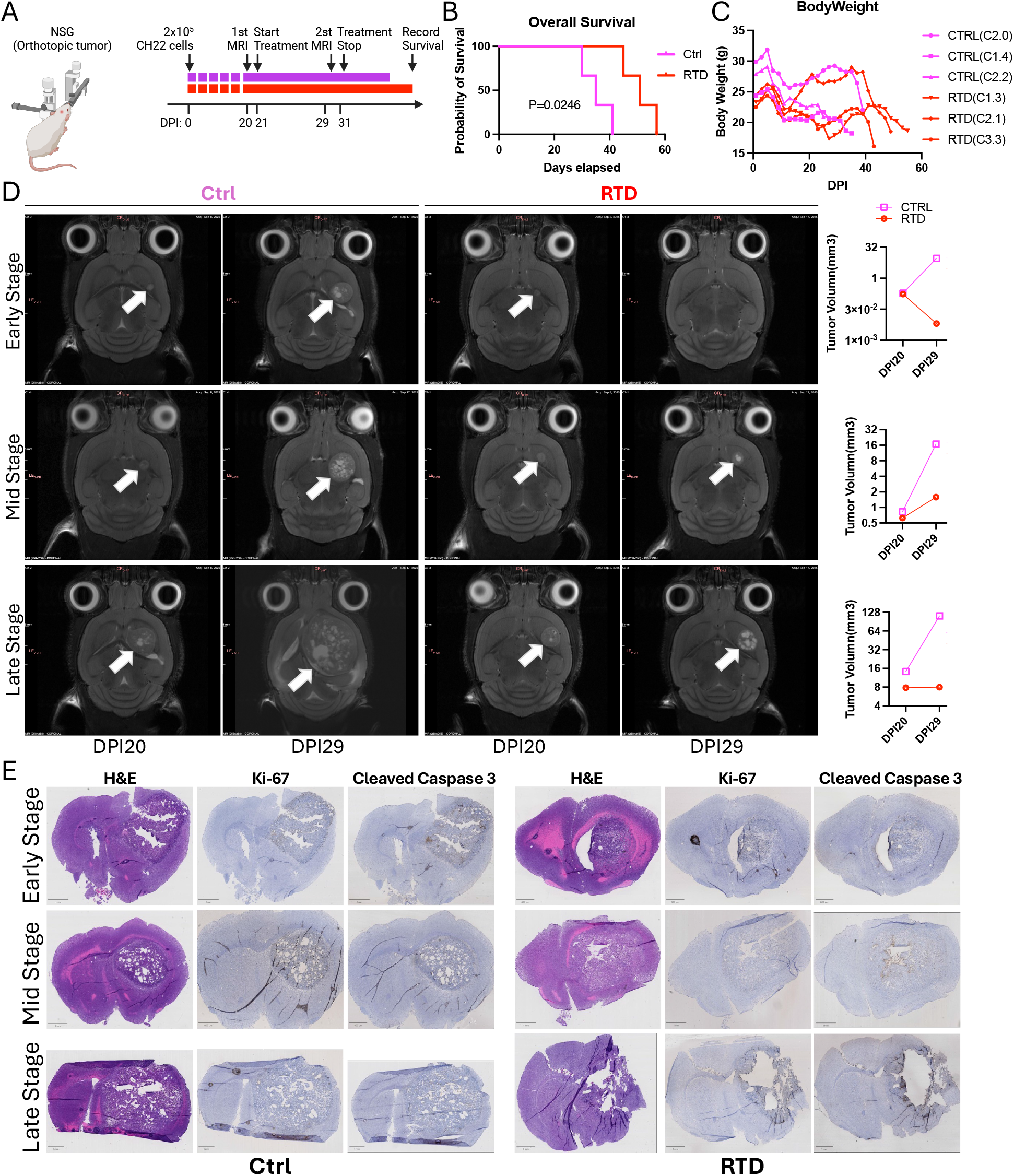
FASN inhibition with low-dose fractionated RT suppresses orthotopic chordoma growth. (A) Schematic of the orthotopic chordoma model, dosing, and monitoring schedule with two arms: CTRL (vehicle) and RTD (low-dose fractionated radiotherapy, LDFR: 1 Gy/day for 4 days, plus TVB-2640 25 mg/kg/day for 10 days; combination). Each group included 3 mice (n = 3). (B) Kaplan–Meier survival analysis shows that RTD combination therapy (LDFR + TVB-2640) significantly prolongs survival of mice bearing orthotopic chordoma compared with CTRL (log-rank test). (C) Body-weight (gram) tracking shows that mice in the RTD group (LDFR + TVB-2640) experienced a decrease in body weight over the treatment period. (D) Longitudinal MRI at 20 and 29 dpi. By the second MRI (29 dpi), CTRL animals—across early-, mid-, and late-stage tumors—showed interval growth. In contrast, RTD-treated mice exhibited no obvious growth in mid/late-stage tumors and developed intratumoral T2-weighted hyperintensity consistent with liquefactive necrosis; in early-stage cases, small tumors became non-visible. (E) H&E with matched IHC for Ki-67 and cleaved caspase-3. RTD tumors show expanded cleaved caspase-3–positive necrotic regions and reduced Ki-67–positive proliferative zones, whereas CTRL exhibits the opposite pattern (scale bar indicated).

## DISCUSSION

Collectively, our result support a unifying model in which radiation therapy (RT) resistance in chordoma is shaped by an adaptive fatty acid metabolic program that buffers oxidative injury and increases survival of RT-resistant, stem-like tumor subpopulations.

Our data found a relatively higher PUFA/MUFA ratio (Figs. 1H, 1I) in primary versus recurrent SBC from patients. PUFAs can be liberated from PUFA-containing phospholipids under oxidative stress and undergo lipid peroxidation, which damages cell membranes leading to cell death^31^. RT induces oxidative stress and a dose-dependent increase in PUFAs (Fig. 3J). In contrast, MUFAs buffer oxidative stress because they are intrinsically less peroxidizable and they can support formation of lipid droplet (LDs), MUFAs are enriched downstream of FASN-driven lipogenesis and MUFA-biased lipid remodeling would be expected to raise the threshold for radiation-induced lipid peroxidation and oxidative injury, creating cell subpopulations better equipped to withstand RT-induced oxidative stress.

The critical point we uncovered is that recurrent SBC showed increased fatty acid biosynthesis, leading to accumulation of both PUFAs and MUFAs. A shift from high to low PUFA/MUFA ratios in recurrent SBC is consistent with increased oxidative stress-buffering by LDs and MUFAs to limit membrane damage and increase survival. During recurrence, there is a redistribution of fatty acids toward more protected storage pools (MUFA-containing LDs) rather than membrane-associated PUFAs, which are less vulnerable to oxidative stress^32^.

Radiation rapidly generates ROS and ER stress responses and activation of the PERK–SREBP1–FASN axis promotes fatty acid biosynthesis. With PUFAs increasing at early stage after RT and MUFAs rising later, suggesting that MUFAs accumulate over time after RT and provide a selective advantage that promotes the survival of RT-tolerant persister cells, a notion that is also supported by the appearance of LDs in CH22R, which acquire RT-tolerance over time.

Interestingly, we observed a spatial distinct distribution of PUFA, which are largely enriched at the proliferative rim, exclusive of MUFAs, in a patient-derived SBC xenograft. Thus, the PUFA/MUFA ratio is low in the established tumor core and increases toward the proliferative rim, alongside largely mutually exclusive MUFA-versus PUFA-enriched regions the core may be more therapy refractory than the rim. In this framework, a MUFA-enriched tumor core may represent an oxidative-stress–adapted metabolic state. This spatial perspective suggests that, from the standpoint of fatty acids in RT resistance, it may be time to look beyond the invasive front and pay greater attention to the metabolically resilient tumor core, which may represent a critical niche driving tumor recurrence^33–35^.

Our *in-silico* knockout and pharmacologic inhibition of FASN identified it as a functional vulnerability within RT resistant cells. Pharmacologic FASN inhibition restored RT sensitivity *in vitro*, normalizing levels of DNA damage in RT-tolerant cells and significantly blocking tumor growth in combination with low-dose RT in a subcutaneous as well as orthotopic patient-derived SBC xenograft model. FASN catalyzes a rate-limiting step in lipogenesis, and its overexpression is linked to resistance to genotoxic treatments— including chemo- and radiotherapy—across multiple cancers^25,36–38^. Because small-molecule FASN inhibitors are clinically available, this approach could be integrated with standard-of-care surgery plus RT on SBC and possibly other types of chordoma. Together, these findings support FASN inhibition as a practical radiosensitization strategy for chordoma, particularly in SBC, where dose escalation is constrained by anatomy.

Unlike single-gene targets, metabolic interventions are not strictly tumor-specific and can influence adjacent or systemic tissues^39,40^. In our models the RT + TVB-2640 showed one tail-tip necrosis in the subcutaneous model and one post-treatment alopecia in the intracranial model (Supplement Fig.S5). The side effect profile of systemic FASN inhibitors is limited, and importantly, this lack of exclusivity is not solely a drawback. Highly selective, single-target therapies often falter in the face of intratumoral heterogeneity and adaptive escape. In contrast, targeting an adaptive metabolic phenotype—particularly when combined with established adjuvant modalities—can provide broader coverage across coexisting tumor subpopulations and evolving resistance trajectories. Adjuvant intracranial radiotherapy is a mainstay for SBC but carries high risks for treatment-induced morbidity. Combination strategies such as combining RT with FASN inhibition may offer a practical and near-term path to improve control. More broadly, this framework may also be relevant as a later-line option for other cancers that recur with marked heterogeneity over time^41–43^.

### Limitations of the study

1. PERK has been previously implicated in transmitting ROS stress to SREBP1 activation^44^, However, how PERK engagement induces SREBP1 maturation and nuclear entry in chordoma remains an open question.
2. We have yet to visualize PUFA integration into LD on a subcellular level to show a direct connection between LD expansion and PUFA sequestering and redox buffering.
3. As a rare cancer, chordoma offers limited access to primary tissue, and, as a bone-derived sarcoma, there are few robust cell lines that perform reliably in TC environment for extensive drug and radiotherapy testing. CH22 is the only relatively robust cell line that has been widely used in recent chordoma studies. Next steps will need to incorporate additional patient-derived organoids robust enough for preclinical testing to strengthen biological and translational relevance.

## RESOURCE AVAILABILITY

### Lead contact

Requests for further information and resources should be directed to and will be fulfilled by the lead contact, Claudia Petritsch.

### Materials availability

This study did not generate new unique reagents.

### Data and code availability

RNA-seq and global Nano-ESI mass spectrometry datasets generated in this study will be submitted to journal-designated public repositories and made available upon publication; additional data and code are available from the corresponding author upon reasonable request.

## ACKNOWLEDGMENTS

We thank Dr. Mengcheng Shen for valuable technical advice on Western blot experiments. We thank Chordoma Foundation’s Josh Sommer and Ashley White for generously providing the cell lines and for their additional support and assistance throughout this work. We thank Dr. Jieun Kim of the Neurosciences Preclinical Imaging Lab (NPIL) at the Wu Tsai Neurosciences Institute for technical assistance with MRI of the mouse models.

## AUTHOR CONTRIBUTIONS

R.W., C.P., conceived the ideas. R.W., Y.M., and C.P. designed the experiments. R.W., Y.M., E.N., D.P., and Y.L.X performed the experiments. Y.M. conducted MS. R.W., Y.M., and C.P. analyzed the data. R.W. and H.O. conducted biostatistics and bioinformatics analysis. R.N,Z. provided MS resource. J.F.M provided surgery sample. R.W., Y.M., and C.P. wrote the manuscript, and all authors reviewed and approved the manuscript for publication.

## DECLARATION OF GENERATIVE AI AND AI-ASSISTED TECHNOLOGIES

During the preparation of this work, the authors used ChatGPT (OpenAI) to assist with literature searching and language editing. After using this tool or service, the author(s) reviewed and edited the content as needed, and we take full responsibility for the content of the publication.

## SUPPLEMENTAL INFORMATION

Document S1. Figures S1–S8, Tables S1 and S2

Document S2. All Samples—Nano-ESI MS Full-scan Raw MS1 Spectrum

Document S3. Unprocessed Western blot images (full membranes)

## FUNDING

National Institute of Health R21 NS099836 (C.K.P)

National Institute of Health R01 NS080619 (C.K.P)

National Institute of Health R01 CA164746 (C.K.P.)

National Institute of Health 17X074 (C.K.P.)

Neurosciences Preclinical Imaging Lab Pilot Grant 2022 (C.K.P.)

Stanford Women’s Health and Sex Differences Center (WHSDM) Seed Grant (C.K.P)

BRAF LGG consortium research fund (C.K.P.)

Stanford Cancer Institute (C.K.P)

National Institute of Health U54 CA261717 (C.K.P.)

Department of Neurosurgery Seed Grand (J.F.M, C.K.P.)

## KEY RESOURCES TABLE

| REAGENT OR RESOURCE | SOURCE | IDENTIFIER |
| --- | --- | --- |
| **Antibodies** |  |  |
| Alexa Fluor 488 phalloidin, IF, dil: 1/200 | Thermo Fisher Scientific | Cat#A12379 |
| Alexa Fluor 647, IF, dil: 1:200 | Thermo Fisher Scientific | Cat#A-31571 |
| Anti-mouse IgG, HRP-linked Antibody, WB, dil: 1:5000 | Cell Signaling Technology | Cat#7076 |
| Anti-rabbit IgG, HRP-linked Antibody, WB, dil: 1:5000 | Cell Signaling Technology | Cat#7074 |
| beta Actin HRP (BA3R), WB, dil: 1:5000 | Thermo Fisher Scientific | Cat#MA5-15739-HRP |
| Brachyury, IF, dil: 1:100 | R&D Systems | Cat#AF2085 |
| Cleaved Caspase-3 (Asp175), WB, dil: 1/1000 | Cell Signaling Technology | Cat#9661S |
| Fatty Acid Synthase (C20G5), WB, dil: 1/1000 | Cell Signaling Technology | Cat#3180 |
| GADD45A, WB, dil: 1/5000 | Thermo Fisher Scientific | Cat#PA5-88248 |
| GAPDH HRP (GA1R), WB, dil: 1:5000 | Thermo Fisher Scientific | Cat#MA5-15738-HRP |
| PARP1, IF, dil: 1:200; WB, dil: 1:1000 | Thermo Fisher Scientific | Cat#436400 |
| PERK (C33E10), WB, dil: 1:1000 | Cell Signaling Technology | Cat#3192 |
| Phospho-PERK (Thr982), WB, dil: 1:1000 | Thermo Fisher Scientific | Cat#PA5-40294 |
| SREBP-1 (E9F40), WB, dil: 1/1000 | Cell Signaling Technology | Cat#9587 |
| Phospho-yH2AX, IF, dil: 1:200 | Thermo Fisher Scientific | Cat#MA5-33062 |
| **Bacterial and virus strains** |  |  |
| n/a |  |  |
| **Biological samples** |  |  |
| Human Chordoma Samples | Dr.Juan Fernandez Miranda, Department of Neurosurgery, Stanford University |  |
| Human Chordoma Samples FFPE Slides | Department of Neuro-Pathology, Stanford University |  |
| **Chemicals, peptides, and recombinant proteins** |  |  |
| 2-mercaptoethanol, 25mL | BIO-RAD | Cat#1610710 |
| AntiFade Mounting Medium with DAPI | Thermo Fisher Scientific | Cat#NC1848444 |
| BODIPY 493/503, IF, dil: 5uM | MedChemExpress | Cat#HY-W090090 |
| BODIPY 576/589, IF, dil: 5uM | MedChemExpress | Cat#HY-D1118 |
| Bovine Serum Albumin Solution | Millipore Sigma | Cat#A8412 |
| Cell-Titer-Glo | Promega | Cat#G7572 |
| CellROX Green Reagent, IF, dil: 5uM/L | Thermo Fisher Scientific | Cat#C10444 |
| Cytiva HyClone™ Phosphate Buffered Saline (PBS) | Thermo Fisher Scientific | Cat#SH30256LS |
| Denifanstat (TVB-2640) | MedChemExpress | Cat#HY-112829 |
| Desmosterol | MedChemExpress | Cat#HY-113224 |
| EveryBlot Blocking Buffer | BIO-RAD | Cat#12010020 |
| FBS | Gibco | Cat#16000044 |
| GlutaMAX | Gibco | Cat#35050061 |
| IMDM | Gibco | Cat#12440079 |
| Live Cell Imaging Solution | Thermo Fisher Scientific | Cat#A59688DJ |
| MEM NEAA | Corning | Cat#25-025-CIR |
| NuPAGE 4-12% Bis-Tris Gel | Thermo Fisher Scientific | Cat#NP0321BOX |
| Oleic acid | Sigma-Aldrich | Cat#01008-5G |
| Pierce™ BCA Protein Assay Kits | Thermo Fisher Scientific | Cat#23227 |
| Pierce™ LDS Sample Buffer | Thermo Fisher Scientific | Cat#84788 |
| Protease/Phosphatase Inhibitor Cocktail | Cell Signaling Technology | Cat#5872 |
| RIPA Lysis and Extraction Buffer | Thermo Fisher Scientific | Cat#89900 |
| RPMI1640 | Gibco | Cat#11875085 |
| Sodium Chloride 0.9% | Thermo Fisher Scientific | Cat#Z1376 |
| Spectra Multicolor Broad Range Protein Ladder | Thermo Fisher Scientific | Cat#26634 |
| Staurosporine | MedChemExpress | Cat#HY-15141 |
| SuperSignal™ West Dura Extended Duration Substrate | Thermo Fisher Scientific | Cat#34076 |
| Trans-Blot Turbo Mini 0.2 µm PVDF Transfer Packs | BIO-RAD | Cat#1704156 |
| Tris Buffered Saline, powder, pH 8.0 | Millipore Sigma | Cat#T6664 |
| TrypLE™ Express Enzyme (1X) | Thermo Fisher Scientific | Cat#12605036 |
| Tunicamycin | Cell Signaling Technology | Cat#12819 |
| Western Blot Strip-It Biffer | Advanta | Cat#R-03722-C20 |
| **Critical commercial assays** |  |  |
| n/a |  |  |
| **Deposited data** |  |  |
| 13 Chordoma Bulk RNA Sequencing Data | Deposited at journal-designated repository upon publication |  |
| 13 Chordoma Non-Target Nano-ESI-MS Sequencing Data | Deposited at journal-designated repository upon publication |  |
| **Experimental models: Cell lines** |  |  |
| Human Chordoma Cell Line: CH22 | Chordoma Foundation | CH22 |
| Human Chordoma Cell Line: U-CH11 | Chordoma Foundation | U-CH11 |
| **Experimental models: Organisms/strains** |  |  |
| Athymic nude (Nu/J) mice | Jackson Laboratory (Jax) | 002019 |
| NSG mice | Stanford Veterinary Service Center (VSC) | https://med.stanford.edu/vsc.html |
| **Oligonucleotides** |  |  |
| n/a |  |  |
| **Recombinant DNA** |  |  |
| n/a |  |  |
| **Software and algorithms** |  |  |
| Image J | NIH | Version 2.16.0 |
| Image Lab | BIO-RAD Laboratories Inc. | Version 6.1.0 build 7 |
| Prism | GraphPad Software, San Diego, CA | Version 10 |
| Qupath | QuPath developers, The University of Edinburgh | Version 0.6.0 |
| R | The R Foundation for Statistical Computing | Version 4.5.1 |
| R Studio | Posit Software, PBC | Version 2025.05.1+513 |
| Weasis | Weasis Team | Version 4.6.3 |
| **Other** |  |  |
| n/a |  |  |

## EXPERIMENTAL MODEL AND STUDY PARTICIPANT DETAILS

### Cell culture and generation of stable cell lines

Cell culturing follows recommendation protocol from Chordoma Foundation. Specifically, CH22 U-CH11 cells were cultured in a mix of IMDM and RPMI-1640 with 4:1 ratio, supplemented with 10% FBS, 1% NEAA and 1% GlutaMAX, at 37 °C in a humidified atmosphere of 95% air and 5% CO2. Details of the CH22-derived cell variants established through different treatment conditions are provided in Table S1.

### Mice

Athymic nude (Nu/J) mice with 6–8 weeks of age were obtained from Jackson Laboratory (Jax) and acclimated for at least 2 days to generate chordoma subcutaneous xenograft models. NSG mice with 6–8 weeks of age were obtained from established in-house breeding colony to generate chordoma orthotopic xenograft models. Mice were housed 5 maximum per cage in a conventional barrier facility on a 12-hour light/dark cycle at 22°C with free access to water and food. Mice health status was checked by following the protocols. All animal procedures were performed in accordance with institutional and federal guidelines (Stanford University APLAC protocol # 33633).

### Generation and monitoring of orthotopic glioma models

6-8-week-old NSG mice were used for orthotopic implantation of CH22 chordoma cells. CH22 cells were harvested from in vitro culture during the exponential growth phase, washed twice in sterile phosphate-buffered saline (PBS), and resuspended at a density of 1 ×10^8 cells/mL in 50% Cultrex®/PBS, 2uL per mouse. Mice were anesthetized with isoflurane inhalation (4% induction, 2% maintenance) and ice block, respectively. The head of the anesthetized mouse was fixed in a stereotaxic frame using a nose cone and ear bars. Eyes were moistened with water-based lubricant, and the fur was cleared over the head with an electric shaver. A sterile cotton tip soaked in betadine was used to swab and clean the surface of the incision site. A midline incision was made on the scalp, and a small hole was drilled into the skull to expose the brain surface at the following stereotaxic coordinates selected to target the striatum, relative to bregma (anterior-posterior–1.00 mm, medial-lateral–1.80 mm). Cells were gently drawn up into a blunt-tipped 26G needle attached to a 10μL syringe (Hamilton Company, Reno, NV). The syringe was then placed into a microinjection pump (UltraMicroPump III, World Precision Instruments, Sarasota, FL) attached to the stereotaxic frame and lowered slowly into the injection site at a depth of 3 mm for adult mice and 1 mm for postnatal mice. The microinjection pump controlled the injection of cells at a flow rate of 0.5 μL/min, after which the needle was left in place for 5 min to ensure complete diffusion of cells and avoid backflow. Following the slow withdrawal of the needle, the hole was closed with dental cement, the incision was sutured, and the mice were observed until full recovery from anesthesia on a heated pad.

Mice were monitored every other day. Animals assessed for survival and body weight recording were monitored until they lost more than 20% of their original body weight and/or had clinical symptoms of tumor development (hydrocephalus, lethargy, hunched posture).

### Generation and monitoring of subcutaneous chordoma models

6-8-week-old Nu/J mice were used for subcutaneous implantation of CH22 chordoma cells. CH22 cells were harvested from in vitro culture during the exponential growth phase, washed twice in sterile phosphate-buffered saline (PBS), and resuspended at a density of 1 ×10^7 cells/mL in 50% Cultrex®/PBS, 100uL per mouse. For subcutaneous tumor implantation, each mouse was immobilized under brief isoflurane sedative. A 26-gauge needle was used to inject 100 µL of the cell suspension into the right flank subcutaneous space. After injection, mice were monitored until fully recovered from sedative and returned to their home cage.

Mice were monitored every other day. Tumor growth was recorded by caliper measurement of two perpendicular diameters (length and width). Tumor volume (V) was calculated using the formula:

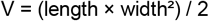

All monitoring procedures and humane endpoints were established in accordance with institutional IACUC protocols and the OBSERVE: guidelines for the refinement of rodent cancer models, ensuring minimal pain and distress. Mice were euthanized by CO_2_ asphyxiation followed by cervical dislocation when tumors reached 1,000 mm^3^, or showed signs of ulceration, >10% body-weight loss, or impaired mobility.

## METHOD DETAILS

### Mice Magnetic resonance imaging

All MR studies were performed at 7T Bruker BioSpec system (Bruker, Billeria, MA) equipped with BGA-12S gradient insert and interfaced to ParaVision 360 V. 3.5. A 4-element receive-only mouse CryoProbe and 86mm volume coil were used for MR imaging. Animals were anesthetized with 2.5% isoflurane mixed in air and O2 and maintained with 1–1.5% isoflurane during the imaging procedure. Core body temperature was maintained at 37°C using a warm water circulator pad and a temperature controller. The animal’s respiration was also monitored via a pneumatic pad placed under the animal (SA Instruments, Inc., Stony Brook, NY). T2w Turbo RARE sequence was used to obtain coronal (TR/TE = 2000/30 ms, FOV = 20 × 20 mm2, matrix = 256 × 256, slice thickness = 0.5 mm, RARE Factor = 4, NEX = 2, scan time = 4 min 16 s) and transverse (TR/TE = 2500/30 ms, FOV = 20 × 20 mm^2^, matrix = 256 × 256, slice thickness = 0.5 mm, RARE Factor = 4, NEX = 2, scan time = 5 min 20 s) structural MRI.

Mice bearing orthotopic tumors were imaged on day 20 post inoculation (DPI 20) and again on day 29 (DPI 29). The transverse and coronal slice showing the largest cross-sectional area of the tumor was identified, and three orthogonal tumor diameters—length (L), width (W), and height (H)—were measured from consecutive transverse and coronal images using open-source software Weasis (v4.6.3 https://github.com/nroduit/Weasis). Tumor volume (V) was calculated according to the ellipsoid formula:

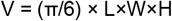

where V is in mm^3^ (with L, W and H in mm).

### Tissue collection and processing of mouse samples

Mice were intraperitoneally injected with ketamine (100 mg/kg) and xylazine (10 mg/kg) to induce deep anesthesia, and subsequently transcardiac perfusion was performed. For mice designated for DESI-MS imaging, perfusion was performed with sterile deionized (DI) water only. Brains were then excised, gently rinsed with DI water, and embedded in 4% carboxymethyl cellulose (CMC) for cryosectioning. For other mice, perfusion was performed first with phosphate-buffered saline followed by 4% paraformaldehyde (PFA)/PBS. Brains perfused with 4% PFA were then resected, post-fixed in 4% PFA/PBS for 2 hr on ice, cryoprotected in 15% sucrose/PBS for 24 hr and then 30% sucrose/PBS for at least 48 hr, followed by embedding in Tissue-Tek O.C.T. Compound (Sakura Finetek USA, Inc., Torrance, CA). The embedded tissues were stored at −80°C until sectioned. 10μm-thick coronal sections of the brain were cut on a cryostat (Leica, Wetzlar, Germany), collected onto Superfrost Plus slides (Menzel Glaser, VWR) and air dried for 1h before storing at −80°C until stained. H&E, Ki67 and Cleaved-Caspase3 staining were using Histo-Tech protocols (Histo-Tec Laboratory Inc., Hayward, CA; http://histoteclab.com/).

### Human studies

Tumor specimens were obtained from patients undergoing surgical resection at the Department of Neurosurgery, Stanford Healthcare. Written informed consent was obtained from all patients in accordance with protocols approved by the Institutional Review Board (IRB) of Stanford University. Immediately after surgical removal, tumor tissues were placed on ice in sterile phosphate-buffered saline (PBS) and transported to the laboratory within 30 minutes. Necrotic and non-tumor areas were trimmed away, and viable tumor regions were snap-frozen in liquid nitrogen and stored at −80 °C for subsequent bulk-RNA and metabolomic sequencing and analyses.

For each case, demographic and clinical variables were abstracted from de-identified medical record, including age, sex, anatomic tumor site, and primary versus recurrent status. The final chordoma diagnosis for all cases was rendered by board-certified pathologists in the Stanford Pathology Department, integrating histomorphology and immunohistochemistry (IHC). Consistent with current practice, brachyury (TBXT) IHC was used to support the diagnosis and distinguish chordoma from histologic mimics. Formalin-fixed, paraffin-embedded (FFPE) tissue section slides were retrieved from the Department of Pathology; H&E and TBXT (brachyury) IHC slides used for review and downstream analyses were issued by the pathology service.

### Western Blot

Cells that were cultured in 6-well plates were washed three times with cold phosphate-buffered saline (PBS) and lysed directly in each well using 100 µL of RIPA lysis buffer (Thermo Fisher, Cat# 89900) supplemented with 1% protease and phosphatase inhibitor cocktail (100×, Thermo Fisher). Lysates were scraped, transferred to 1.5 mL microtubes, and stored at −80 °C until use. For protein extraction, thawed samples were sonicated on ice using a Bioruptor Pico (Diagenode) and centrifuged at 12,000 × g for 10 min at 4 °C. The supernatant was collected, and protein concentration was quantified using the bicinchoninic acid (BCA) assay (Thermo Fisher, Cat# 23225) following the manufacturer’s instructions. Equal amounts of protein (30 µg per sample) were mixed with 4× LDS loading buffer (Invitrogen, Cat# NP0007) containing 5% β-mercaptoethanol and denatured at 95 °C for 5 min. Samples were separated by SDS-PAGE using 4–12% Bis-Tris precast gels (Invitrogen, Cat# NP0322) in 1× MOPS running buffer and subsequently transferred onto PVDF membranes (Bio-Rad, Cat# 162-0177) using the Trans-Blot Turbo system (Bio-Rad; 25 V, 1.3 A, 7 min). Membranes were blocked for 5 min with commercial fast-blocking buffer (Advansta) or with 3% bovine serum albumin (BSA) in Tris-buffered saline containing 0.1% Tween-20 (TBST), followed by incubation with primary antibodies diluted in 3% BSA/TBST overnight at 4 °C with gentle agitation. After three washes in TBST (5 min each), membranes were incubated with HRP-conjugated secondary antibodies (1:5,000 in TBST) for 1 hr at room temperature and washed again. Protein bands were visualized using enhanced chemiluminescence (ECL) substrate (SuperSignal West Femto, Thermo Fisher) and imaged with a ChemiDoc imaging system (Bio-Rad) under high-sensitivity mode. When required, membranes were stripped using stripping buffer (Advansta, Cat# R-03722-D50; 5 mL per blot, 37 °C for 45 min) and reprobed. Total protein staining was performed with Pierce Reversible Protein Stain (Thermo Fisher) according to the manufacturer’s protocol for normalization and loading control verification.

### Immunocytochemistry (ICC) and confocal microscopy

Cells grown on chamber slides (MatTek CCs-8) were fixed with 4% paraformaldehyde (PFA) for 10 min at room temperature (RT), followed by three washes with phosphate-buffered saline (PBS, 5 min each). Fixed cells were permeabilized with 0.5% Triton X-100 in PBS for 20 min at RT and then washed again three times with PBS. Nonspecific binding was blocked for 1hr at RT using blocking buffer (4% BSA and 0.1% Tween-20 in PBS). Cells were then incubated with primary antibodies diluted in blocking buffer overnight at 4 °C in a humidified chamber. The next day, slides were washed three times in PBS (5 min each) and incubated with appropriate fluorophore-conjugated secondary antibodies (1:500 in blocking buffer) for 1 hr at RT in the dark. After three final PBS washes, the chambers were removed, and coverslips were mounted using DAPI-containing antifade mounting medium (Vector Laboratories, Cat# H-1200). Edges were sealed with clear nail polish and allowed to dry for at least 30 min before imaging.

Confocal fluorescence images were acquired using a Zeiss LSM 780 confocal microscope equipped with 405 nm, 488 nm, 561 nm, and 633 nm laser lines. Imaging parameters were kept constant across experimental groups, and representative fields were captured at 20× air objective and 63× oil immersion objective. All images were processed using Fiji (ImageJ, NIH) for brightness/contrast adjustment and channel merging without nonlinear manipulation. Quantification of fluorescence intensity was performed using ImageJ built-in measurement tools, analyses were performed in a blinded fashion.

### Bulk RNA sequencing and transcriptomic analysis

Fresh-frozen tumor specimens stored at −80 °C were sent to Novogene (Novogene Corporation Inc., Sacramento, CA) for bulk RNA sequencing. Samples with RNA integrity number (RIN) ≥ 2.5 were used for library preparation and sequenced on an Illumina NovaSeq 6000 platform to generate 150-bp paired-end reads. Downstream analyses were performed in R (version 4.5.1) with R Studio (Version 2025.05.1 Build513). Genes with an adjusted p < 0.05 and |log_2_ fold change| > 1 were considered differentially expressed genes (DEGs) using the DESeq2 package. Data visualization and exploratory analyses were performed in R using ggplot2, pheatmap, and EnhancedVolcano packages. Volcano plots were generated to display DEGs. Hierarchical clustering and heatmaps of normalized expression values (variance-stabilizing transformation, VST) were used to illustrate expression patterns across samples. Functional enrichment analysis of DEGs was conducted using clusterProfiler for Gene Ontology (GO) and Kyoto Encyclopedia of Genes and Genomes (KEGG) pathway annotation. Significantly enriched pathways were defined as p.adjust < 0.05. The results were visualized as barplots and dot plots. All analyses were performed using custom R scripts, and all figures were generated with consistent color palettes for publication.

### Single cell data acquire and analysis

Open-access single-cell transcriptomic data was obtained from Zhang, Qilin et al from NGDC-GSA database (BioProject No. PRJCA009785, DAC NO.HDAC001399)^45^. Downstream analyses were performed using established single-cell computational workflows on R/R Studio^46^. Briefly, count matrices and accompanying metadata were imported into Seurat and subjected to quality control, normalization, identification of highly variable features, scaling, and graph-based unsupervised clustering with low-dimensional visualization by PCA/UMAP (and t-SNE where indicated). Cell-type annotation was performed using SingleR with reference atlases from celldex, and annotations were further refined by canonical marker inspection. To reconstruct cellular state trajectories, pseudotime analyses were conducted using Monocle (DDRTree). To interrogate gene-level causality, we implemented in silico single-cell perturbation (virtual knockout) analyses using scTenifoldKnk, followed by pathway-level interpretation where appropriate. In parallel, metabolic pathway activities were quantified at single-cell resolution using scMetabolism (VISION/KEGG) to generate module scores that were compared across clusters and conditions. Differential analyses and functional enrichment were carried out using standard statistical tests with multiple-testing correction, and all figures were generated in R using reproducible scripts (e.g., ggplot2, patchwork, pheatmap).

### Nano-ESI-mass spectrometry and data analysis

Fresh-frozen tumor specimens (stored at −80 °C) were subjected to metabolite extraction prior to nano-ESI-MS. On ice, we prepared extraction buffer of 80% methanol/20% acetonitrile (LC–MS grade). We combined1 mg tissue with 50 µL buffer (1 mg:50 µL) and finely minced and homogenized the sample, which was processed through three cycles of 10 s vortex → 20 s on ice → 30 s bath sonication. Samples were then agitated at 4 °C for 30 min, followed by protein precipitation at −20 °C for 2 hr. After centrifugation (12,000 rpm, 10 min, 4 °C), the clarified supernatant was transferred to 2 mL protein low-bind tubes and stored at −80 °C until MS acquisition.

Spectra were acquired in negative ion modes on a Fusion Orbitrap mass spectrometer (Thermo Fisher, Waltham, MA) equipped with a home-built nano-ESI source, using a spray voltage of 1.2 kV, capillary temperature of 200 °C, and full-scan mass range of *m/z* 100–1000 at a resolution of 60,000. Instrumental parameters and sample injection volumes were kept constant across all runs.

Raw MS files (*.RAW) were converted to mzML format using ProteoWizard MSConvert (version 3.0) with centroiding enabled. All downstream processing and statistical analyses were performed in R (version 4.x) using the xcms, MSnbase, and tidyverse packages. Peak detection and retention-time alignment were carried out using the centWave algorithm (ppm = 10, peak width = c(5, 30), snthresh = 10). After peak grouping and missing-value imputation, data matrices were log_2_-transformed and normalized to total ion current (TIC). Multivariate analyses including principal component analysis (PCA) and hierarchical clustering were performed to assess sample separation. Differentially abundant metabolites (DAMs) between experimental groups were identified using Student’s t-test or moderated statistics (p < 0.05, |log_2_ fold change| > 1). Visualization was generated using ggplot2, pheatmap, and EnhancedVolcano to produce volcano plots and heatmaps. Annotation of significant *m/z* features was performed by matching accurate mass (tolerance ± 5 ppm) against public databases, including HMDB, LipidMaps, and Metlin. All nano-ESI-MS data processing pipelines and visualization scripts were implemented in R with fully reproducible code. Representative figures were generated in R using unified color palettes consistent with data visualization.

### DESI-mass spectrometry and data visualization

Cells grown on glass coverslips and tissue sections (10 µm thickness) were stored at −80 °C until analysis. Sections were brought to room temperature under desiccation and then transferred to the DESI (desorption electrospray ionization) imaging platform. MSI experiments were carried out with a commercial DESI source (Prosolia Inc., Indianapolis, IN, USA). A commercial DESI sprayer (Viktor Technology Co., Ltd., Beijing, China) was used in this experiment. The inner diameter (i.d.) of the DESI spray capillary was 20 μm, and the outer diameter (o.d.) of the capillary was 120 μm. During the imaging process, the Y-distance between the DESI tip and the sample surface was set as 4 mm. The X-distance between the DESI tip and the MS inlet was set as 2 mm. The impact angle between the DESI sprayer and the sample stage was set at 60°. Compressed N_2_ (120 psi, 99.999% purity) was used as the nebulizing gas. A negative voltage (−6 kV) was applied to the DESI sprayer. All mass spectra were recorded in negative ion mode. The injection rate of DESI solvent (ACN:DMF = 1:1) was set as 0.7 μL/min. During the data-collecting process, the moving speed of the X-axis of the sample platform was set as 300 μM/s, while the step size of the Y-axis was set as 100 μm. The spatial resolution of DESI-MSI is about 100 microns. An Orbitrap mass spectrometer (Fusion, Thermo Fisher Scientific, Waltham, MA, USA) was used to obtain MS data. Negative full MS scans were employed over the range of *m/z* 200– 1000. The automatic gain control (AGC) was set to the off position to keep the scan rate constant. The mass resolution was set to 30,000. The maximum injection time was set as 100 μS. The MS inlet capillary temperature was set at 300 °C. MS images are generated using a commercial software (MassImager Pro, Chemmind Technology, Beijing, China).

## QUANTIFICATION AND STATISTICAL ANALYSIS

All significance calculations were conducted using GraphPad Prism software. Statistical significance was determined using an unpaired, two-tailed Student’s t test or two-way ANOVA method when more than two groups were being compared. Survival was analyzed by the Kaplan-Meier method using log rank (Mantel-Cox) test. Quantitative data are reported as mean ± SEM, and any values of p < 0.05 were considered statistically significant. ∗p < 0.05, ∗∗p < 0.01, ∗∗∗p < 0.001 and ∗∗∗∗p < 0.0001.

## REFERENCES

1. Jones, P.S., Aghi, M.K., Muzikansky, A., Shih, H.A., Barker, F.G., and Curry, W.T. (2014). Outcomes and patterns of care in adult skull base chordomas from the Surveillance, Epidemiology, and End Results (SEER) database. J Clin Neurosci 21, 1490–1496. 10.1016/j.jocn.2014.02.008.

2. McMaster, M.L., Goldstein, A.M., Bromley, C.M., Ishibe, N., and Parry, D.M. (2001). Chordoma: incidence and survival patterns in the United States, 1973-1995. Cancer Causes Control 12, 1–11. 10.1023/a:1008947301735.

3. Sharifnia, T., Hong, A.L., Painter, C.A., and Boehm, J.S. (2017). Emerging Opportunities for Target Discovery in Rare Cancers. Cell Chem Biol 24, 1075–1091. 10.1016/j.chembiol.2017.08.002.

4. Stacchiotti, S., Sommer, J., and Chordoma Global Consensus Group (2015). Building a global consensus approach to chordoma: a position paper from the medical and patient community. Lancet Oncol 16, e71–83. 10.1016/S1470-2045(14)71190-8.

5. Salisbury, J.R. (1993). The pathology of the human notochord. J Pathol 171, 253–255. 10.1002/path.1711710404.

6. George, B., Bresson, D., Herman, P., and Froelich, S. (2015). Chordomas: A Review. Neurosurg Clin N Am 26, 437–452. 10.1016/j.nec.2015.03.012.

7. Halvorsen, S.C., Benita, Y., Hopton, M., Hoppe, B., Gunnlaugsson, H.O., Korgaonkar, P., Vanderburg, C.R., Nielsen, G.P., Trepanowski, N., Cheah, J.H., et al. (2023). Transcriptional Profiling Supports the Notochordal Origin of Chordoma and Its Dependence on a TGFB1-TBXT Network. Am J Pathol 193, 532–547. 10.1016/j.ajpath.2023.01.014.

8. Caneve, P., Schraps, N., Möller, K., Büyücek, S., Lutz, F., Chirico, V., Viehweger, F., Reiswich, V., von Bargen, C., Kind, S., et al. (2025). Brachyury expression is highly specific for chordoma: A tissue microarray study involving 14,976 cancers from 135 different tumor types and subtypes. Ann Diagn Pathol 76, 152448. 10.1016/j.anndiagpath.2025.152448.

9. Vujovic, S., Henderson, S., Presneau, N., Odell, E., Jacques, T., Tirabosco, R., Boshoff, C., and Flanagan, A. (2006). Brachyury, a crucial regulator of notochordal development, is a novel biomarker for chordomas. The Journal of Pathology 209, 157–165. 10.1002/path.1969.

10. Meng, T., Jin, J., Jiang, C., Huang, R., Yin, H., Song, D., and Cheng, L. (2019). Molecular Targeted Therapy in the Treatment of Chordoma: A Systematic Review. Front Oncol 9, 30. 10.3389/fonc.2019.00030.

11. Fiorentino, A., Gregucci, F., Desideri, I., Fiore, M., Marino, L., Errico, A., Di Rito, A., Borghetti, P., Franco, P., Greto, D., et al. (2021). Radiation treatment for adult rare cancers: Oldest and newest indication. Critical Reviews in Oncology/Hematology 159, 103228. 10.1016/j.critrevonc.2021.103228.

12. McDonald, M.W., Linton, O.R., Moore, M.G., Ting, J.Y., Cohen-Gadol, A.A., and Shah, M.V. (2016). Influence of Residual Tumor Volume and Radiation Dose Coverage in Outcomes for Clival Chordoma. Int J Radiat Oncol Biol Phys 95, 304–311. 10.1016/j.ijrobp.2015.08.011.

13. Perez-Vega, C., Akinduro, O.O., Ruiz-Garcia, H.J., Ghaith, A.K.A., Almeida, J.P., Jentoft, M.E., Mahajan, A., Janus, J.R., Bendok, B.R., Choby, G.W., et al. (2024). Extent of Surgical Resection as a Predictor of Tumor Progression in Skull Base Chordomas: A Multicenter Volumetric Analysis. World Neurosurg 181, e620–e627. 10.1016/j.wneu.2023.10.101.

14. Listenberger, L.L., Han, X., Lewis, S.E., Cases, S., Farese, R.V., Ory, D.S., and Schaffer, J.E. (2003). Triglyceride accumulation protects against fatty acid-induced lipotoxicity. Proceedings of the National Academy of Sciences 100, 3077–3082. 10.1073/pnas.0630588100.

15. Gehrmann, W., Würdemann, W., Plötz, T., Jörns, A., Lenzen, S., and Elsner, M. (2015). Antagonism Between Saturated and Unsaturated Fatty Acids in ROS Mediated Lipotoxicity in Rat Insulin-Producing Cells. Cell Physiol Biochem 36, 852–865. 10.1159/000430261.

16. Nakajima, S., Gotoh, M., Fukasawa, K., Murakami-Murofushi, K., and Kunugi, H. (2019). Oleic acid is a potent inducer for lipid droplet accumulation through its esterification to glycerol by diacylglycerol acyltransferase in primary cortical astrocytes. Brain Res 1725, 146484. 10.1016/j.brainres.2019.146484.

17. Mei, S., Ni, H.-M., Manley, S., Bockus, A., Kassel, K.M., Luyendyk, J.P., Copple, B.L., and Ding, W.-X. (2011). Differential roles of unsaturated and saturated fatty acids on autophagy and apoptosis in hepatocytes. J Pharmacol Exp Ther 339, 487–498. 10.1124/jpet.111.184341.

18. Lombardi, S., Goldman, A.R., Tang, H.-Y., Kossenkov, A.V., Liu, H., Zhou, W., Herlyn, M., Lin, J., and Zhang, R. (2023). Targeting Fatty Acid Reprogramming Suppresses CARM1-expressing Ovarian Cancer. Cancer Res Commun 3, 1067–1077. 10.1158/2767-9764.CRC-23-0030.

19. Mason, P., Liang, B., Li, L., Fremgen, T., Murphy, E., Quinn, A., Madden, S.L., Biemann, H.-P., Wang, B., Cohen, A., et al. (2012). SCD1 Inhibition Causes Cancer Cell Death by Depleting Mono-Unsaturated Fatty Acids. PLOS ONE 7, e33823. 10.1371/journal.pone.0033823.

20. Magtanong, L., Ko, P.-J., To, M., Cao, J.Y., Forcina, G.C., Tarangelo, A., Ward, C.C., Cho, K.Y., Patti, G.J., Nomura, D.K., et al. (2019). Exogenous Monounsaturated Fatty Acids Promote a Ferroptosis-Resistant Cell State. Cell Chem Biol 26, 420-432.e9. 10.1016/j.chembiol.2018.11.016.

21. Doll, S., Proneth, B., Tyurina, Y.Y., Panzilius, E., Kobayashi, S., Ingold, I., Irmler, M., Beckers, J., Aichler, M., Walch, A., et al. (2017). Acsl4 Dictates Ferroptosis Sensitivity by Shaping Cellular Lipid Composition. Nat Chem Biol 13, 91–98. 10.1038/nchembio.2239.

22. Azzam, E.I., Jay-Gerin, J.-P., and Pain, D. (2012). Ionizing radiation-induced metabolic oxidative stress and prolonged cell injury. Cancer Lett 327, 48–60. 10.1016/j.canlet.2011.12.012.

23. Cheng, X., Li, J., and Guo, D. (2018). SCAP/SREBPs are Central Players in Lipid Metabolism and Novel Metabolic Targets in Cancer Therapy. Curr Top Med Chem 18, 484–493. 10.2174/1568026618666180523104541.

24. Sánchez-Álvarez, M., Lolo, F.N., Sailem, H., Fulgoni, G., Pascual-Vargas, P., Agüera, L., Catalá-Montoro, M., Arias-García, M., López, J.A., Vázquez, J., et al. (2025). PERK-dependent reciprocal crosstalk between ER and non-centrosomal microtubules coordinates ER architecture and cell shape. Cell Rep 44, 115590. 10.1016/j.celrep.2025.115590.

25. Chen, J., Zhang, F., Ren, X., Wang, Y., Huang, W., Zhang, J., and Cui, Y. (2020). Targeting fatty acid synthase sensitizes human nasopharyngeal carcinoma cells to radiation via downregulating frizzled class receptor 10. Cancer Biol Med 17, 740–752. 10.20892/j.issn.2095-3941.2020.0219.

26. Zhan, N., Li, B., Xu, X., Xu, J., and Hu, S. (2018). Inhibition of FASN expression enhances radiosensitivity in human non-small cell lung cancer. Oncol Lett 15, 4578–4584. 10.3892/ol.2018.7896.

27. Chuang, H.-Y., Lee, Y.-P., Lin, W.-C., Lin, Y.-H., and Hwang, J.-J. (2019). Fatty Acid Inhibition Sensitizes Androgen-Dependent and -Independent Prostate Cancer to Radiotherapy via FASN/NF-κB Pathway. Sci Rep 9, 13284. 10.1038/s41598-019-49486-2.

28. Ye, L.F., Chaudhary, K.R., Zandkarimi, F., Harken, A.D., Kinslow, C.J., Upadhyayula, P.S., Dovas, A., Higgins, D.M., Tan, H., Zhang, Y., et al. (2020). Radiation-Induced Lipid Peroxidation Triggers Ferroptosis and Synergizes with Ferroptosis Inducers. ACS Chem Biol 15, 469–484. 10.1021/acschembio.9b00939.

29. O’Farrell, M., Duke, G., Crowley, R., Buckley, D., Martins, E.B., Bhattacharya, D., Friedman, S.L., and Kemble, G. (2022). FASN inhibition targets multiple drivers of NASH by reducing steatosis, inflammation and fibrosis in preclinical models. Sci Rep 12, 15661. 10.1038/s41598-022-19459-z.

30. Wang, X., Du, Q., Mai, Q., Zou, Q., Wang, S., Lin, X., Chen, Q., Wei, M., Chi, C., Peng, Z., et al. (2025). Targeting FASN enhances cisplatin sensitivity via SLC7A11-mediated ferroptosis in cervical cancer. Translational Oncology 56, 102396. 10.1016/j.tranon.2025.102396.

31. Zhang, S., Zhang, J., Fan, X., Liu, H., Zhu, M., Yang, M., Zhang, X., Zhang, H., and Yu, F. (2022). Ionizing Radiation-Induced Ferroptosis Based on Nanomaterials. Int J Nanomedicine 17, 3497–3507. 10.2147/IJN.S372947.

32. Qiu, B., Ackerman, D., Sanchez, D.J., Li, B., Ochocki, J.D., Grazioli, A., Bobrovnikova-Marjon, E., Diehl, J.A., Keith, B., and Simon, M.C. (2015). HIF2α-Dependent Lipid Storage Promotes Endoplasmic Reticulum Homeostasis in Clear-Cell Renal Cell Carcinoma. Cancer Discov 5, 652–667. 10.1158/2159-8290.CD-14-1507.

33. Buglakova, E., Ekelöf, M., Schwaiger-Haber, M., Schlicker, L., Molenaar, M.R., Shahraz, M., Stuart, L., Eisenbarth, A., Hilsenstein, V., Patti, G.J., et al. (2024). Spatial single-cell isotope tracing reveals heterogeneity of de novo fatty acid synthesis in cancer. Nat Metab 6, 1695–1711. 10.1038/s42255-024-01118-4.

34. Cao, Y., Li, J., Chen, Y., Wang, Y., Liu, Z., Huang, L., Liu, B., Feng, Y., Yao, S., Zhou, L., et al. (2025). Monounsaturated fatty acids promote cancer radioresistance by inhibiting ferroptosis through ACSL3. Cell Death Dis 16, 184. 10.1038/s41419-025-07516-0.

35. Lei, G., Zhuang, L., and Gan, B. (2024). The roles of ferroptosis in cancer: Tumor suppression, tumor microenvironment, and therapeutic interventions. Cancer Cell 42, 513–534. 10.1016/j.ccell.2024.03.011.

36. Jin, Y., Chen, Z., Dong, J., Wang, B., Fan, S., Yang, X., and Cui, M. (2021). SREBP1/FASN/cholesterol axis facilitates radioresistance in colorectal cancer. FEBS Open Bio 11, 1343–1352. 10.1002/2211-5463.13137.

37. Yang, L., Zhang, F., Wang, X., Tsai, Y., Chuang, K.-H., Keng, P.C., Lee, S.O., and Chen, Y. (2016). A FASN-TGF-β1-FASN regulatory loop contributes to high EMT/metastatic potential of cisplatin-resistant non-small cell lung cancer. Oncotarget 7, 55543–55554. 10.18632/oncotarget.10837.

38. Tadros, S., Shukla, S.K., King, R.J., Gunda, V., Vernucci, E., Abrego, J., Chaika, N.V., Yu, F., Lazenby, A.J., Berim, L., et al. (2017). De Novo Lipid Synthesis Facilitates Gemcitabine Resistance through Endoplasmic Reticulum Stress in Pancreatic Cancer. Cancer Res 77, 5503–5517. 10.1158/0008-5472.CAN-16-3062.

39. Poon, D.J.J., Tay, L.M., Ho, D., Chua, M.L.K., Chow, E.K.-H., and Yeo, E.L.L. (2021). Improving the therapeutic ratio of radiotherapy against radioresistant cancers: Leveraging on novel artificial intelligence-based approaches for drug combination discovery. Cancer Letters 511, 56–67. 10.1016/j.canlet.2021.04.019.

40. van Gisbergen, M.W., Zwilling, E., and Dubois, L.J. (2021). Metabolic Rewiring in Radiation Oncology Toward Improving the Therapeutic Ratio. Front Oncol 11, 653621. 10.3389/fonc.2021.653621.

41. Rojas, M., Manzi, M., Madurga, S., García Velásquez, F.E., Romero, M.A., Marín, S., Cascante, M., and Maurel, J. (2025). Metabolic plasticity drives specific mechanisms of chemotherapy and targeted therapy resistance in metastatic colorectal cancer. Explor Target Antitumor Ther 6, 1002337. 10.37349/etat.2025.1002337.

42. Yu, Y., Yu, J., Ge, S., Su, Y., and Fan, X. (2023). Novel insight into metabolic reprogrammming in cancer radioresistance: A promising therapeutic target in radiotherapy. Int J Biol Sci 19, 811–828. 10.7150/ijbs.79928.

43. Liu, S., Zhang, X., Wang, W., Li, X., Sun, X., Zhao, Y., Wang, Q., Li, Y., Hu, F., and Ren, H. (2024). Metabolic reprogramming and therapeutic resistance in primary and metastatic breast cancer. Mol Cancer 23, 261. 10.1186/s12943-024-02165-x.

44. Bobrovnikova-Marjon, E., Hatzivassiliou, G., Grigoriadou, C., Romero, M., Cavener, D.R., Thompson, C.B., and Diehl, J.A. (2008). PERK-dependent regulation of lipogenesis during mouse mammary gland development and adipocyte differentiation. Proceedings of the National Academy of Sciences 105, 16314–16319. 10.1073/pnas.0808517105.

45. Zhang, Q., Fei, L., Han, R., Huang, R., Wang, Y., Chen, H., Yao, B., Qiao, N., Wang, Z., Ma, Z., et al. (2022). Single-cell transcriptome reveals cellular hierarchies and guides p-EMT-targeted trial in skull base chordoma. Cell Discov 8, 94. 10.1038/s41421-022-00459-2.

46. Wei, R., Xie, H., Zhou, Y., Chen, X., Zhang, L., Bui, B., and Liu, X. (2024). VCAN in the extracellular matrix drives glioma recurrence by enhancing cell proliferation and migration. Front Neurosci 18, 1501906. 10.3389/fnins.2024.1501906.

