## Supplemental File1 for "FASN Inhibition Resensitizes Chordoma to Radiotherapy by Targeting Adaptive Unsaturated Fatty Acid Metabolism"

[1]Stanford University School of Medicine, Stanford, CA 94305, USA.
[2]Department of Neurosurgery, Stanford University School of Medicine, Stanford CA 94305, USA
[3]College of Chemistry, Frontiers Science Centre for New Organic Matter, Nankai University, Tianjin 300071, China
[4]Department of Chemistry, Stanford University, Stanford, CA 94305, USA
[5]Department of Neurology and Neurosciences, Stanford University School of Medicine, Stanford, CA 94305, USA
[6]Department of Neurology, University of Pittsburgh, Pittsburgh, PA 15123, USA
[7]Pittsburgh Institute for Neurodegenerative Disorders, University of Pittsburgh, Pittsburgh, PA, USA
[8]Department of Neurosurgery, Brown University Rhode Island Hospital, Providence, RI 02903, USA
[9]Stanford Cancer Institute, Stanford University School of Medicine, Stanford, CA 94305, USA
[10]Cancer Model Development Center, Stanford University School of Medicine, Stanford, CA 94305, USA
[11]These authors contributed equally
[12]Senior author
[13]Lead contact

### Supplemental Figures and Tables

**TableS1.** Summary of treatment conditions and time points in cell models

| Cell Line ID | Treatment |
|---|---|
| CH22 | Original chordoma cell line acquired from Chordoma Foundation |
| $CH22^R$ | CH22 treated with repeated irradiation of 1Gy/day for 20 days plus 2 weeks/3 consecutive passage |
| $CH22^{OA}$ | CH22 treated with medium with 300 μM oleic acid for 3 consecutive passages |
| R-CH22 | CH22 treated with one-time irradiation |
| R-$CH22^R$ | $CH22^R$ treated with one-time irradiation |
| R-$CH22^{OA}$ | $CH22^{OA}$ treated with one-time irradiation |
| D-CH22 | CH22 treated with TVB2640 |
| R-D-CH22 | CH22 treated with TVB2640 then treated with one-time irradiation |
| R-D-$CH22^R$ | $CH22^R$ treated with TVB2640 then treated with one-time irradiation |
| R-D-$CH22^{OA}$ | $CH22^{OA}$ treated with TVB2640 then treated with one-time irradiation |

**TableS2.** Clinical and analytical metadata for skull base chordoma samples and epilepsy normal brain tissue processed as formalin-fixed paraffin-embedded (FFPE) tissue and profiled by RNA-seq and nano-ESI-MS, including sample ID, patient age, sex, and primary vs recurrent status.

| SampleID | SampleType | PatientAge (yrs) | PatientSex | Primary/Recurrent |
|---|---|---|---|---|
| STN151 | Skull Base Chordoma | 11 | Male | Recurrent |
| STN153 | Skull Base Chordoma | 4 | Male | Primary |
| STN159 | Skull Base Chordoma | 33 | Female | Primary |
| STN160 | Skull Base Chordoma | 68 | Female | Primary |
| STN162 | Skull Base Chordoma | 73 | Female | Recurrent |
| STN172 | Skull Base Chordoma | 49 | Male | Recurrent |
| STN180 | Skull Base Chordoma | 46 | Female | Primary |
| STN181 | Skull Base Chordoma | 36 | Male | Recurrent |
| STN208 | Skull Base Chordoma | 74 | Female | Recurrent |
| STN210 | Skull Base Chordoma | 12 | Male | Primary |

| | | | | |
|---|---|---|---|---|
| STN213 | Skull Base Chordoma | 24 | Male | Primary |
| STN223 | Skull Base Chordoma | 36 | Male | Primary |
| STN230 | Skull Base Chordoma | 42 | Male | Primary |

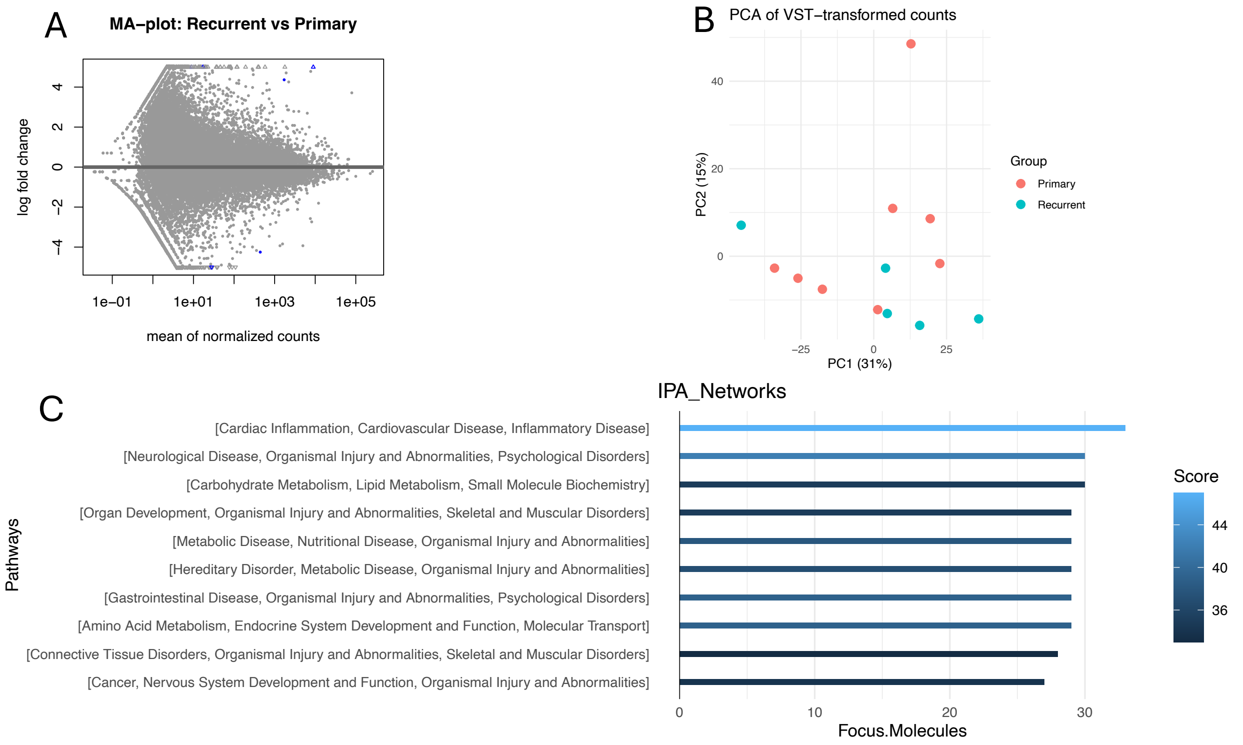


**FigureS1.** (A) MA plot showing log2 fold change of recurrent chordoma versus primary chordoma with mean normalized expression for all genes; each point is a gene, with differentially expressed features highlighted to indicate up- and down-regulation. (B) Principal component analysis (PCA) of normalized transcriptomes separates samples of all chordoma samples, by biological condition along PC1/PC2. (C) Ingenuity Pathway Analysis (IPA) showed that, among the top 10 upregulated networks in recurrent chordoma compared with primary chordoma, metabolic pathways accounted for ~30%, including carbohydrate metabolism, metabolic disease, and amino acid metabolism.

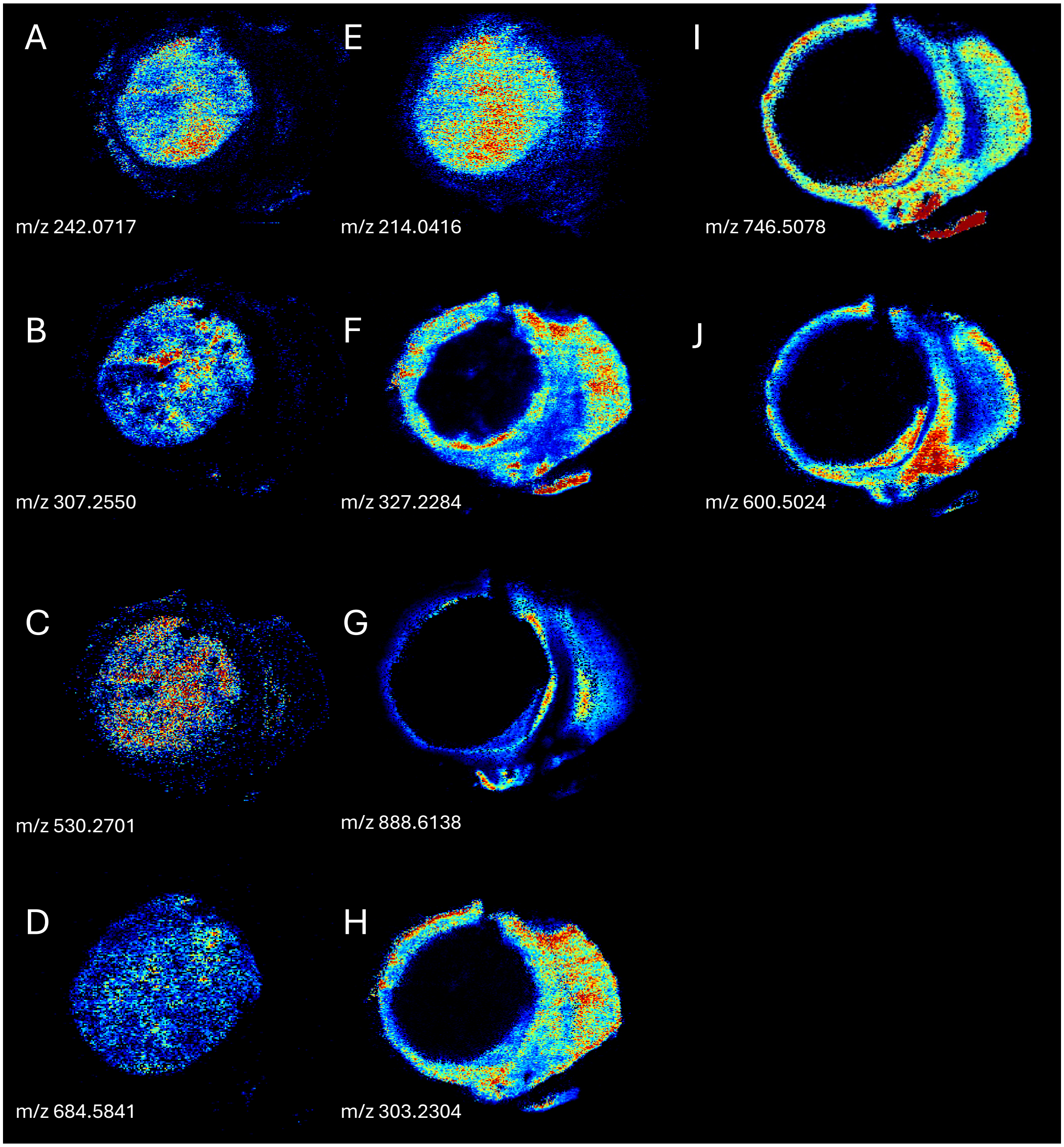


**FigureS2.** (A) DESI–MS image map of NAE 10:2;O2 (m/z 242.0717) on section from mouse orthotopic tumor model. (B) DESI–MS image map of FA 20:2 (m/z 307.2550. (C) DESI–MS image map of ST 27:5;O6;Gly (m/z 530.2701). (D) DESI–MS image map of CAR 33:0;O3 (m/z 684.5841). (E) DESI–MS image map of Phospholipids GTE (m/z 214.0416). (F) DESI–MS image map of PUFA-FA 22:6 (m/z 327.2284). (G) DESI–MS image map of SHexCer 42:2;O2 (m/z 888.6138). (H) DESI–MS image map of PUFA-Arachidonic acid (ARA, m/z 303.2304). (I) DESI–MS image map of LNAPS 33:1 (m/z 746.5078). (J) DESI–MS image map of CAR 31:4 (m/z 600.5024).

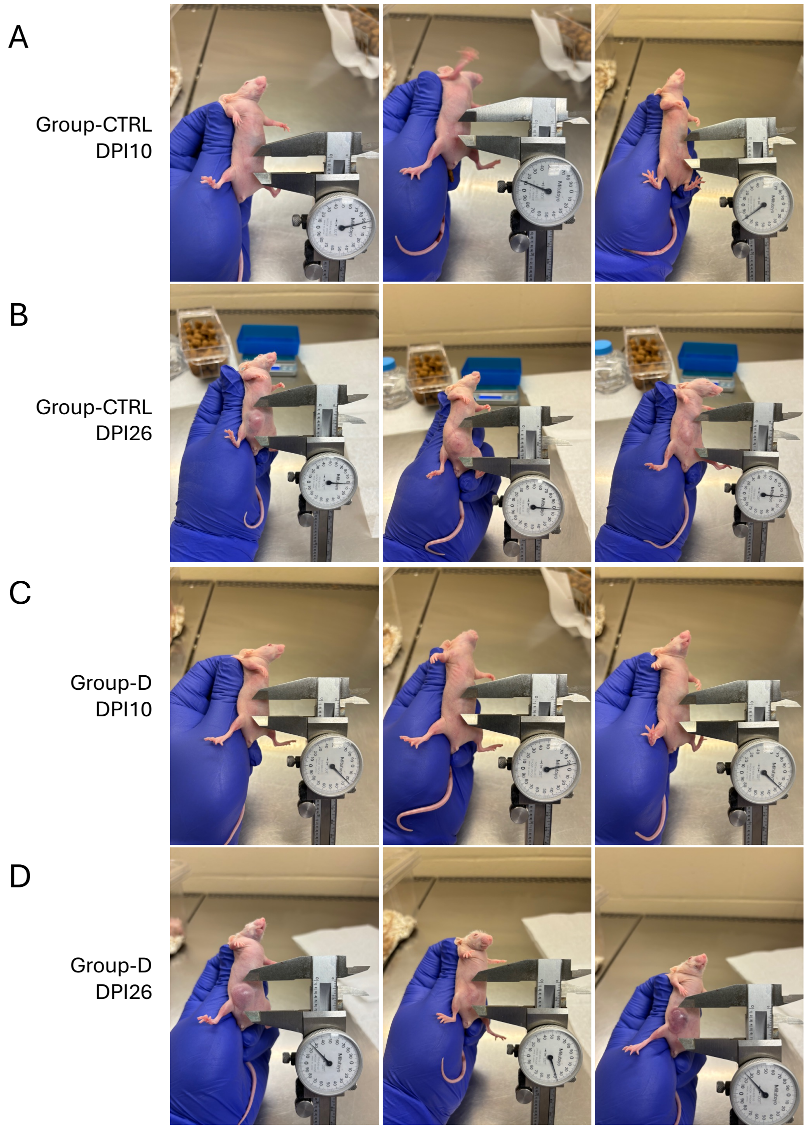


**FigureS3.** (A) Caliper measurement of subcutaneous tumor size in the nude-mouse chordoma xenograft model (Control group, 'Group-CTRL') at 10 days post injection (10 dpi). (B) Caliper measurement of subcutaneous tumor size in the nude-mouse chordoma xenograft model (Control group, 'Group-CTRL') at 26 days post injection (26 dpi). (C) Caliper measurement of subcutaneous tumor size in the nude-mouse chordoma xenograft model (TVB2640 only group, 'Group-D') at 10 days post injection (10 dpi), TVB2640 for 25mg/kg mice body weight/day, 10 days (D) Caliper measurement of subcutaneous tumor size in the nude-mouse chordoma xenograft model (TVB2640 only group, 'Group-D') at 26 days post injection (26 dpi), TVB2640 for 25mg/kg mice body weight/day, 10 days.

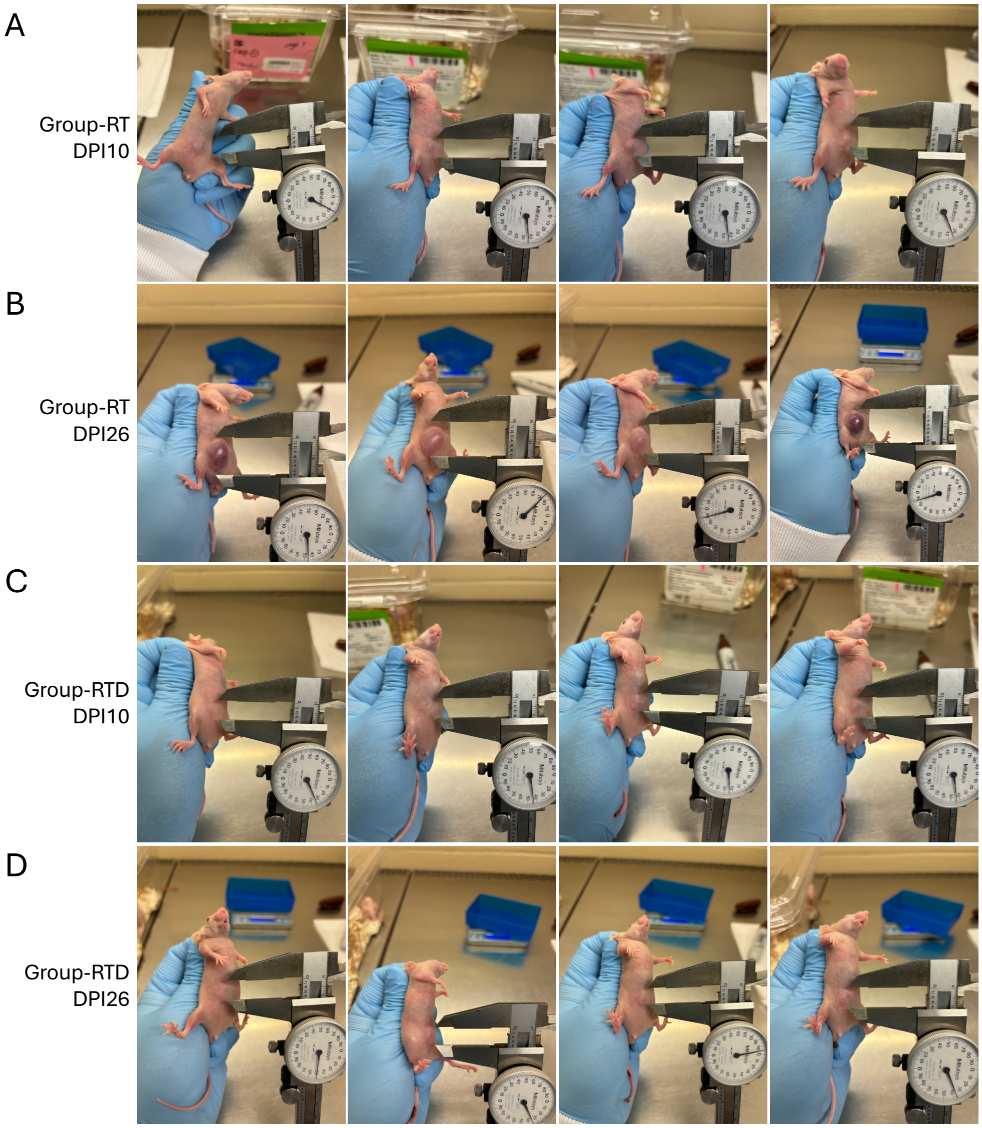


**FigureS4.** (A) Caliper measurement of subcutaneous tumor size in the nude-mouse chordoma xenograft model (Radiotherapy only group, 'Group-RT') at 10 days post injection (10 dpi), radiotherapy using LDFR plan, 1Gy/day, 4days. (B) Caliper measurement of subcutaneous tumor size in the nude-mouse chordoma xenograft model (Radiotherapy only group, 'Group-RT') at 26 days post injection (26 dpi), radiotherapy using LDFR plan, 1Gy/day, 4days. (C) Caliper measurement of subcutaneous tumor size in the nude-mouse chordoma xenograft model (Radiotherapy + TVB2640 group, 'Group-RTD') at 10 days post injection (10 dpi), radiotherapy using LDFR plan, 1Gy/day, 4days. TVB2640 for 25mg/kg mice body weight/day, 10 days. (D) Caliper measurement of subcutaneous tumor size in the nude-mouse chordoma xenograft model (Radiotherapy + TVB2640 group, 'Group-RTD') at 26 days post injection (26 dpi), radiotherapy using LDFR plan, 1Gy/day, 4days. TVB2640 for 25mg/kg mice body weight/day, 10 days.

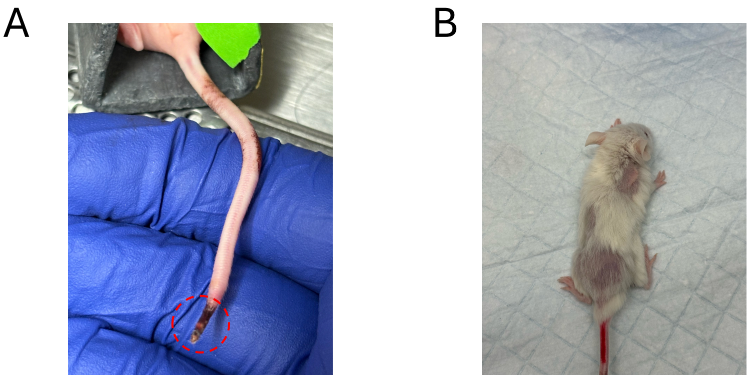

**FigureS5.** (A) One mouse with subcutaneous tumor in the LDFR plus TVB2640 group developed tail necrosis after treatment. (B) One mouse with intracranial tumor in the LDFR plus TVB2640 group developed hair loss after treatment.

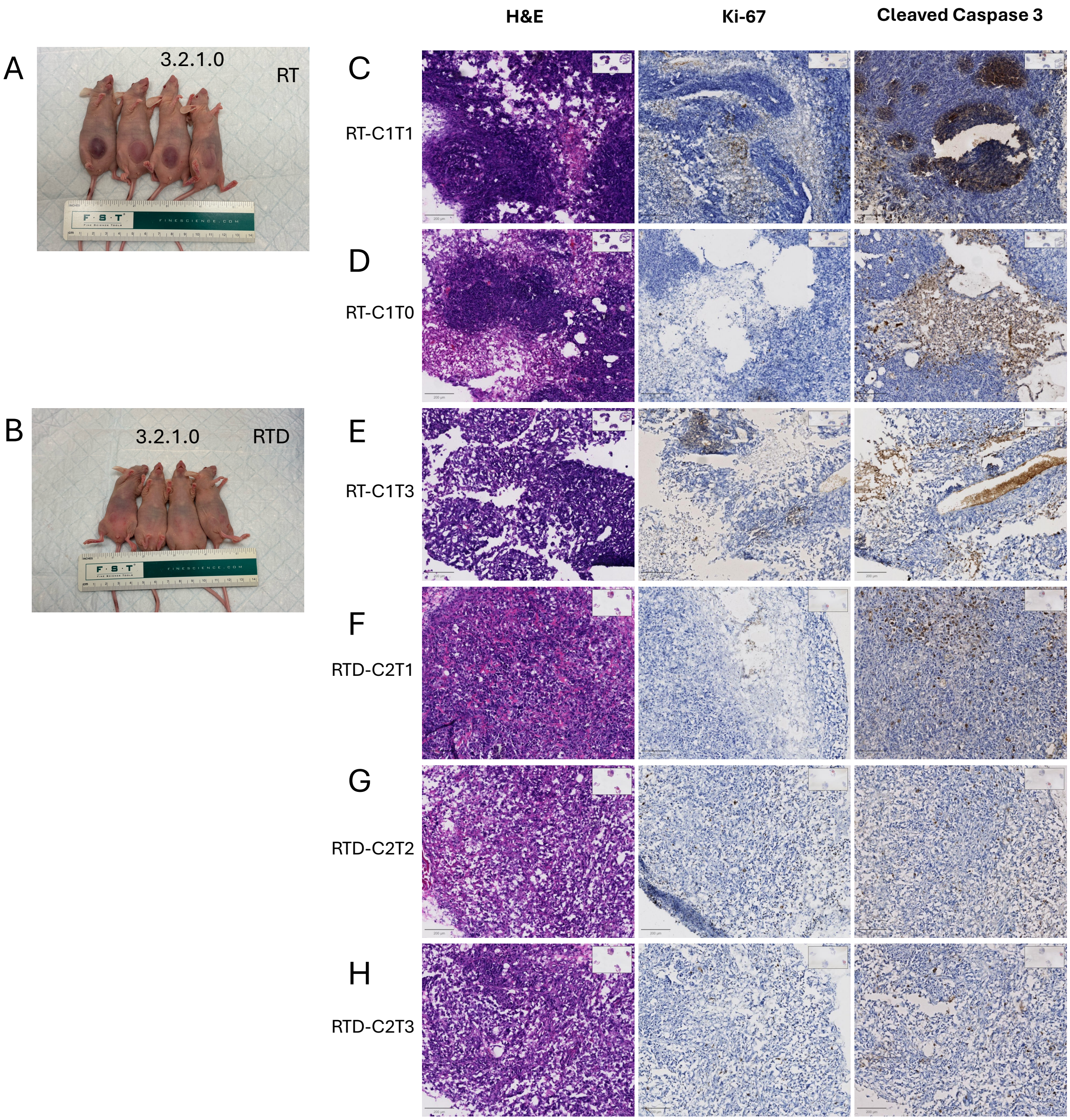


**FigureS6.** (A) Group photograph of Group-RT at the study endpoint showing subcutaneous chordoma tumor burden in nude mice; tumors are visible on the flanks. Ruler in inch and cm for scale. (B) Group photograph of Group-RTD at the study endpoint showing subcutaneous chordoma tumor burden in nude mice; tumors are visible on the flanks. Ruler in inch and cm for scale. (C,D,E) Imaging of H&E and IHC of Ki-67 and Cleaved Caspase 3 of mice from Group-RT. Scale bars = 200µm. (F,G,H) Imaging of H&E and IHC of Ki-67 and Cleaved Caspase 3 of mice from Group-RTD. Scale bars = 200µm.

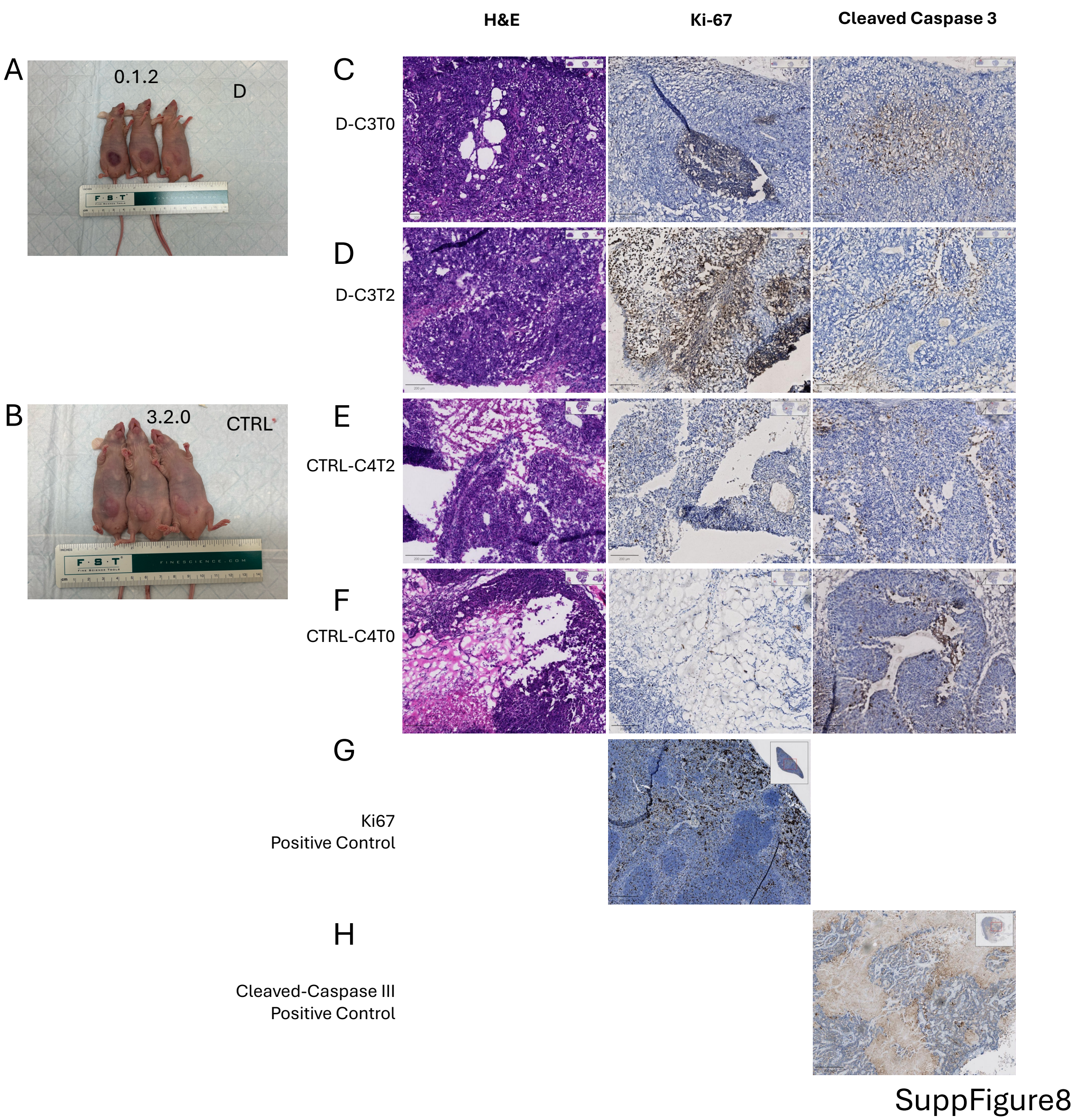


**FigureS7.** (A) Group photograph of Group-D at the study endpoint showing subcutaneous chordoma tumor burden in nude mice; tumors are visible on the flanks. Ruler in inch and cm for scale. (B) Group photograph of Group-CTRL at the study endpoint showing subcutaneous chordoma tumor burden in nude mice; tumors are visible on the flanks. Ruler in inch and cm for scale. (C,D) Imaging of H&E and IHC of Ki-67 and Cleaved Caspase 3 of mice from Group-RT. Scale bars = 200μm. (E,F,) Imaging of H&E and IHC of Ki-67 and Cleaved Caspase 3 of mice from Group-RTD. Scale bars = 200μm. (G) Positive IHC staining control of Ki-67, scale bars = 200μm. (H) Positive IHC staining control of Cleaved Caspase 3, scale bars = 200μm.

**TableS2.** HRMS and MS/MS data or selected ions detected in negative mode Nano-ESI-MS and DESI-MS of brain tissue and chordoma specimens. Measured m/z values were annotated to molecular species on the basis of MS/MS fragmentation, previously reported identifications, or cross-referencing against METABOLOMICS database (https:// https://www.metabolomicsworkbench.org/).

| Source | Measured m/z | Proposed formula | Theoretical m/z | Error(delta) | Attribute |
|---|---|---|---|---|---|
| Nano-ESI-MS | 239.21 | C15H27O2 | 239.2017 | 0.0083 | FA 15:1 |
| Nano-ESI-MS | 267.24 | C17H31O2 | 267.2330 | 0.0070 | FA 17:1 |
| Nano-ESI-MS | 281.26 | C18H33O2 | 281.2486 | 0.0114 | FA 18:1 |
| Nano-ESI-MS | 309.29 | C20H37O2 | 309.2799 | 0.0101 | FA 20:1 |
| Nano-ESI-MS | 365.35 | C24H45O2 | 365.3425 | 0.0075 | FA 24:1 |
| Nano-ESI-MS | 279.24 | C18H31O2 | 279.2330 | 0.0070 | FA 18:2 |
| Nano-ESI-MS | 307.27 | C20H35O2 | 307.2643 | 0.0057 | FA 20:2 |
| Nano-ESI-MS | 305.25 | C20H33O2 | 305.2486 | 0.0014 | FA 20:3 |
| Nano-ESI-MS | 331.26 | C22H35O2 | 331.2643 | 0.0043 | FA 22:4 |
| Nano-ESI-MS | 329.24 | C22H33O2 | 329.2486 | 0.0086 | FA 22:5 |
| DESI-MS | 187.0400 | C5H12O5 | 187.0373 | 0.0027 | Xylitol |
| DESI-MS | 187.0362 | C5H12O5 | 187.0373 | 0.0011 | Xylitol |
| DESI-MS | 303.2304 | C20H32O2 | 303.2330 | 0.0026 | ARA |
| DESI-MS | 281.2472 | C18H33O2 | 281.2486 | 0.0014 | FA 18:1 |
| DESI-MS | 885.5500 | C47H82O13P | 885.5499 | 0.0001 | PI 38:4 |
| DESI-MS | 242.0717 | C12H20NO4 | 242.1398 | 0.0681 | NAE 10:2;O2 |
| DESI-MS | 307.2550 | C20H35O2 | 307.2643 | 0.0093 | FA 20:2 |
| DESI-MS | 530.2701 | C29H40NO8 | 530.2759 | 0.0058 | ST 27:5;O6;Gly |
| DESI-MS | 684.5841 | C40H78NO7 | 684.5784 | 0.0057 | CAR 33:0;O3 |
| DESI-MS | 888.6138 | C48H90NO11S | 888.6206 | 0.0068 | (3'-sulfo)CalCer |
| DESI-MS | 303.2300 | C20H32O2 | 303.2330 | 0.0030 | ARA |
| DESI-MS | 746.5078 | C39H73NO10P | 746.4978 | 0.0100 | LNAPS 33:1 |
| DESI-MS | 600.5024 | C38H66NO4 | 600.4997 | 0.0027 | CAR 31:4 |
| DESI-MS | 214.0416 | C5H14NO6P | 214.0476 | 0.0060 | GTE |
| DESI-MS | 327.2284 | C22H31O2 | 327.2330 | 0.0046 | FA 22:6 |

| | | | | | |
|---|---|---|---|---|---|
| DESI-MS | 834.5219 | C42H76NO15 | 834.5221 | 0.0002 | Hex2Cer 30:2 O4 |
| DESI-MS | 261.0353 | C12H21O6 | 261.1344 | 0.0991 | FA 12:1;O4 |
| DESI-MS | 331.2592 | C22H35O2 | 331.2643 | 0.0051 | FA 22:4 |
| DESI-MS | 279.2226 | C18H31O2 | 279.2330 | 0.0104 | FA 18:2 |
| DESI-MS | 279.2318 | C18H31O2 | 279.2330 | 0.0012 | FA 18:2 |
| DESI-MS | 506.2842 | C24H45NO8P | 506.2888 | 0.0046 | LPE 19:2;O |
| DESI-MS | 305.2474 | C20H33O2 | 305.2486 | 0.0012 | FA 20:3 |
| DESI-MS | 748.5300 | C43H75NO7P | 748.5287 | 0.0013 | PE-O 38:6 |
