## Supplementary figures and images for "FASN Inhibition Resensitizes Chordoma to Radiotherapy by Targeting Adaptive Unsaturated Fatty Acid Metabolism"

### Supplemental File2

#### Supplementary File2:
#### All Samples—Nano-ESI MS Full-scan Raw MS1 Spectrum

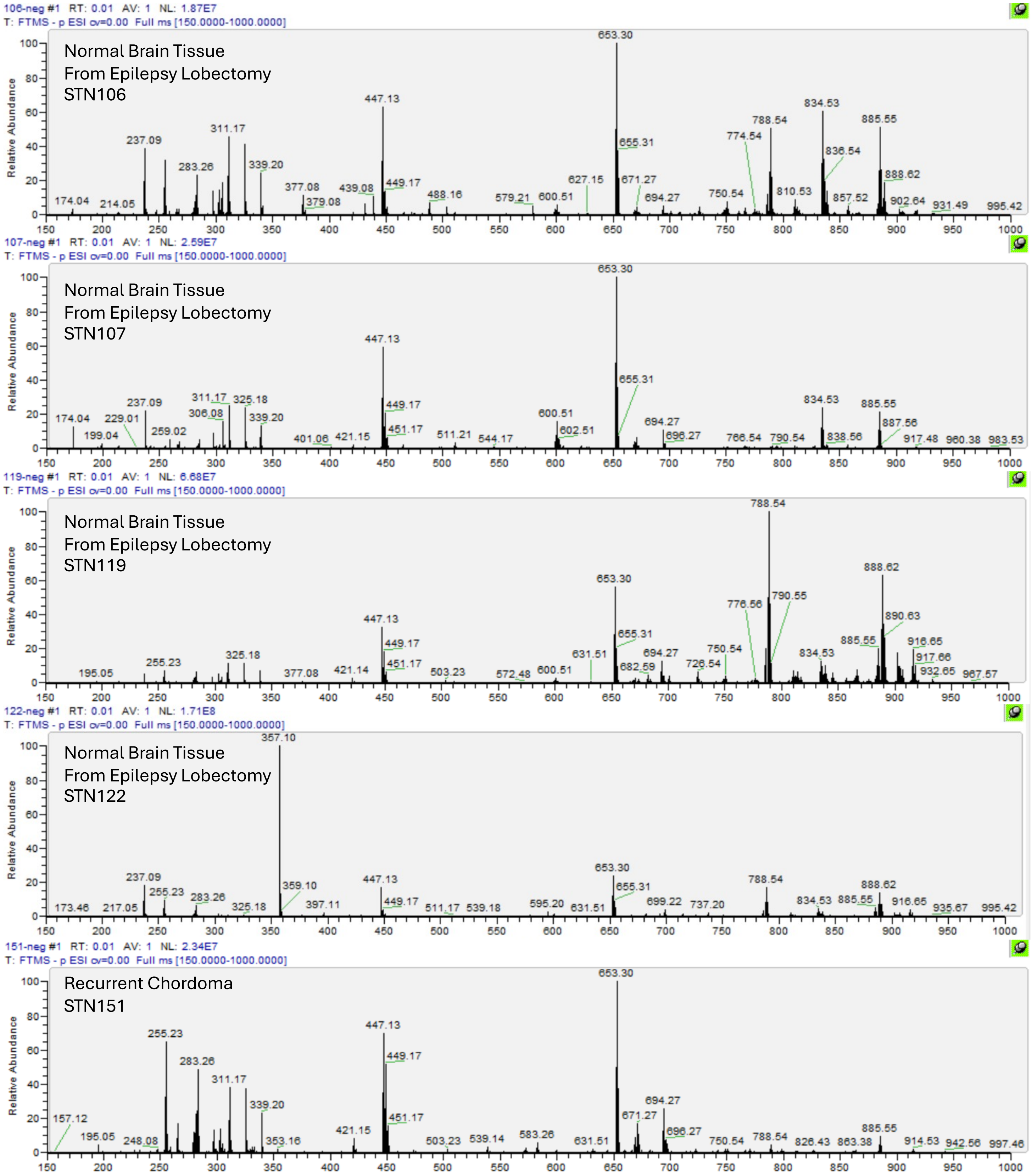

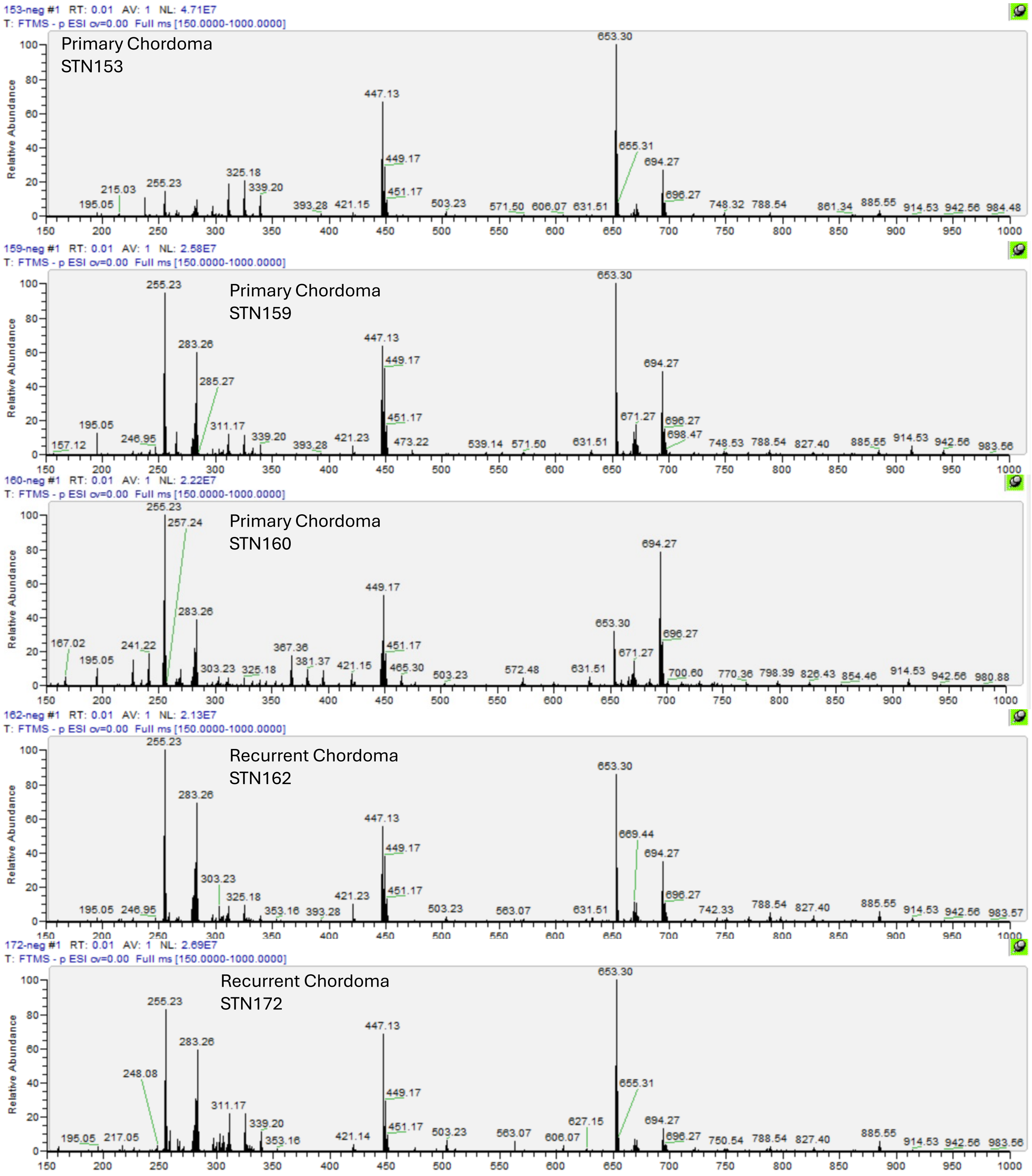

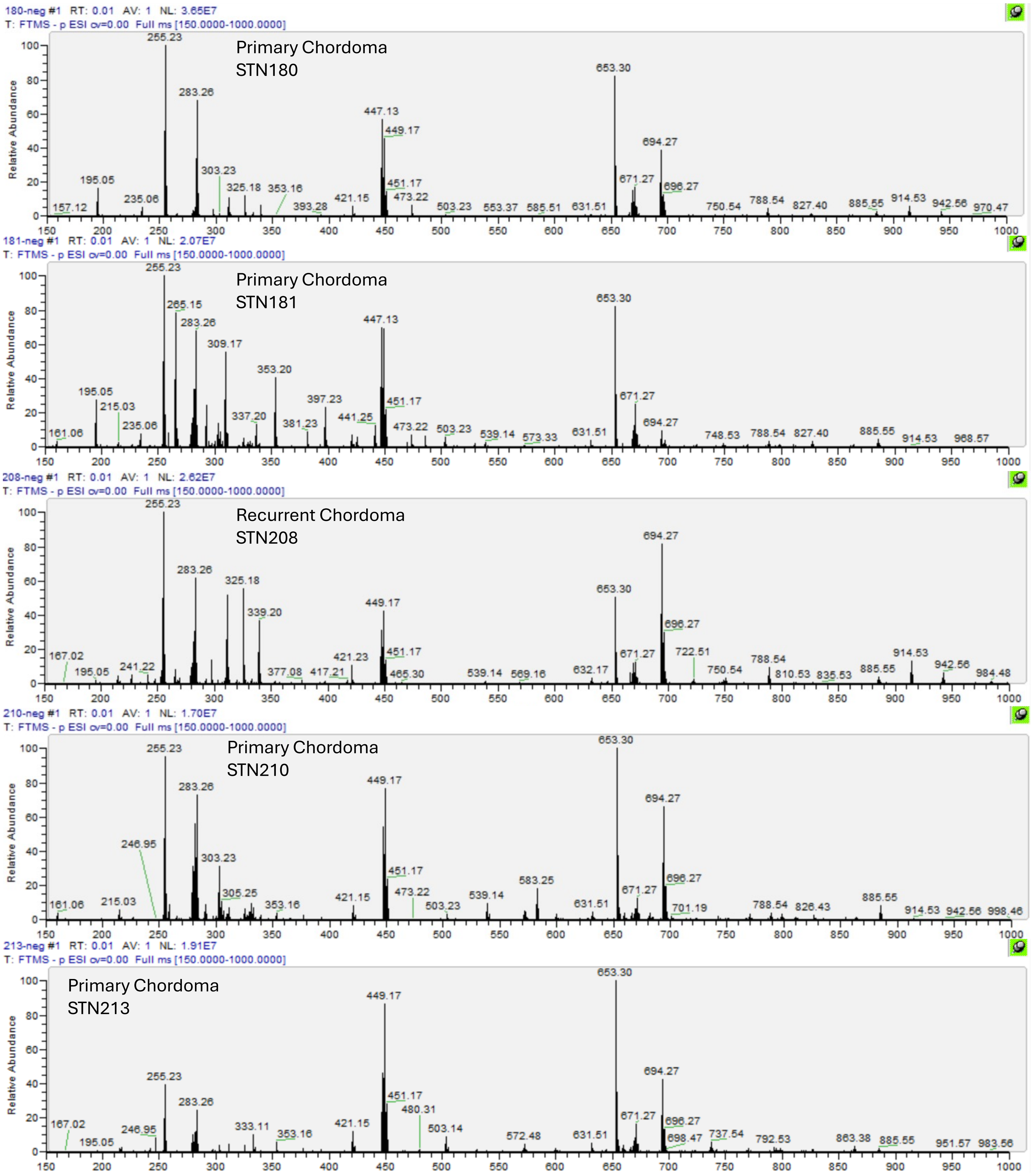

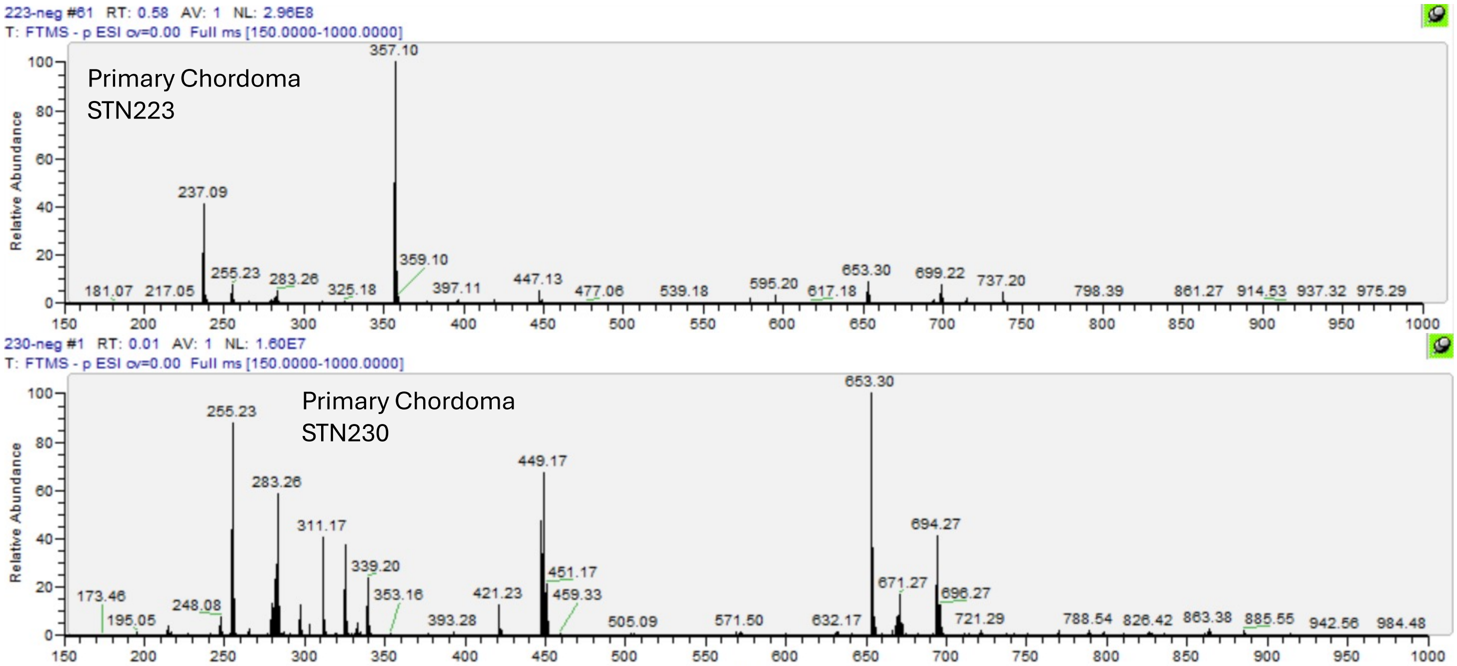
