## Supplemental File3 for "FASN Inhibition Resensitizes Chordoma to Radiotherapy by Targeting Adaptive Unsaturated Fatty Acid Metabolism"

### Supplementary File3:
### Unprocessed Western blot images (full membranes)

Cell: CH22, CH22$^R$
Workflow: FASN-Strip/Reprobe-SREBP1-Strip/Reprobe-PERK-Strip/Reprobe-GAPDH-Memcode

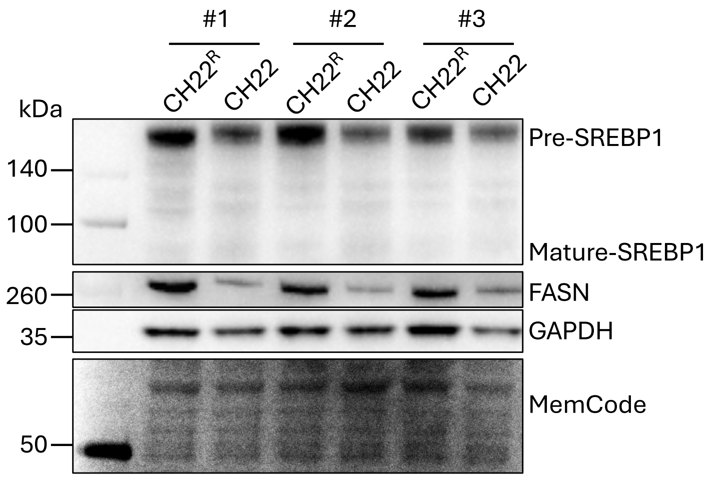


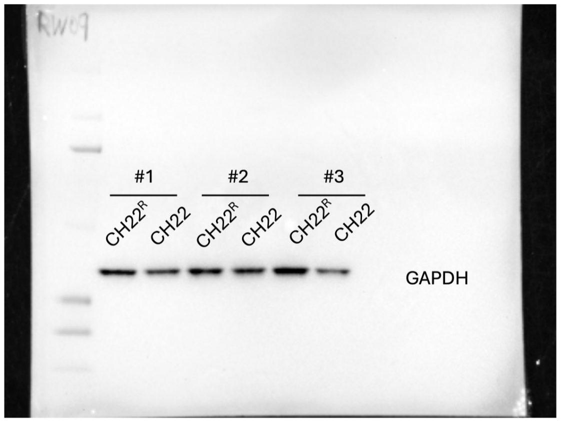


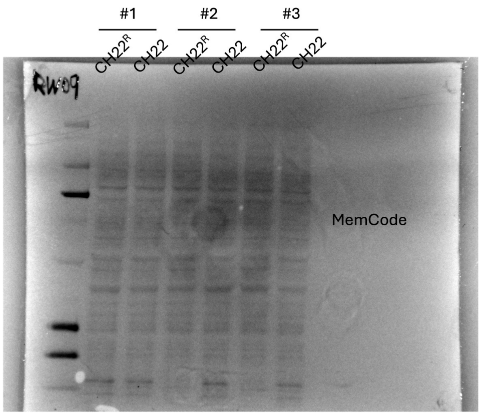


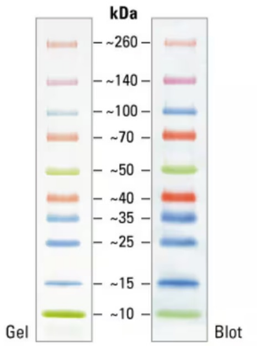


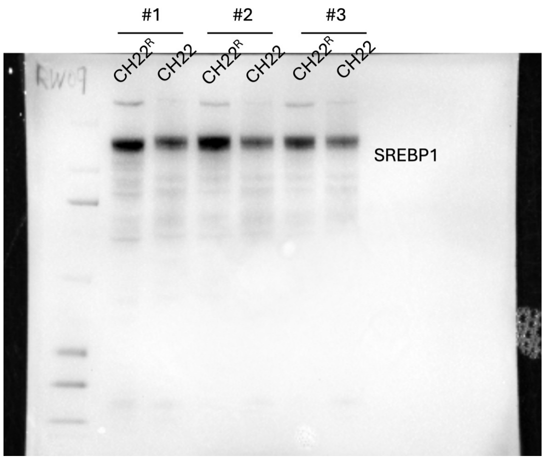


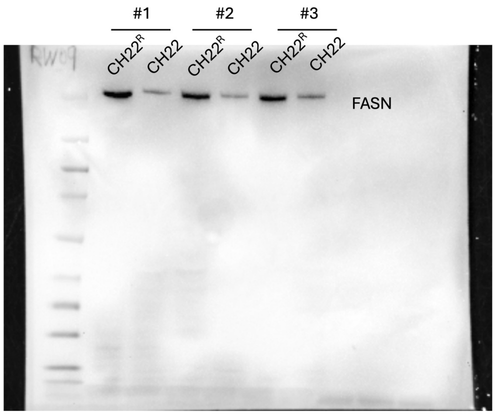

Gel21
Cell: CH22, CH22OA
Workflow: GADD45A-Strip/Reprobe-PARP1-Strip/Reprobe-Cleaved Caspase III-Strip/Reprobe-β Actin-Memcode

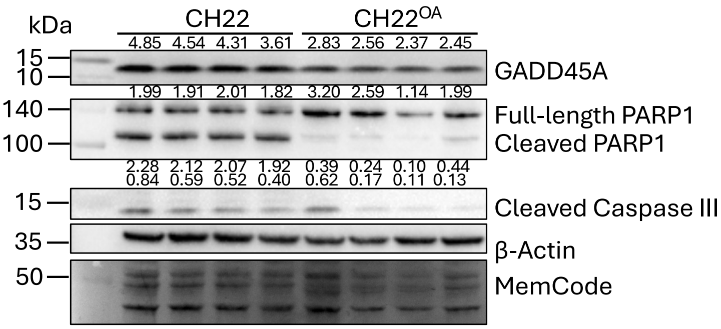


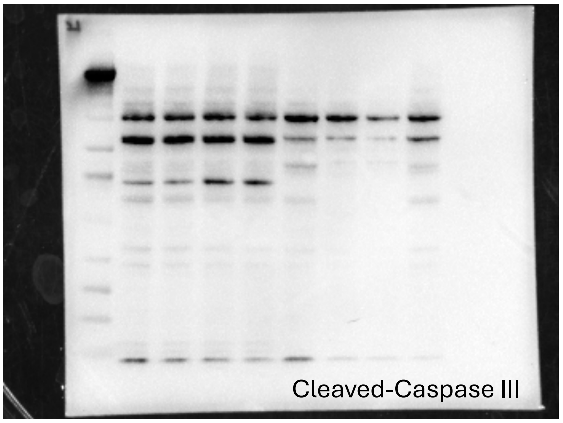


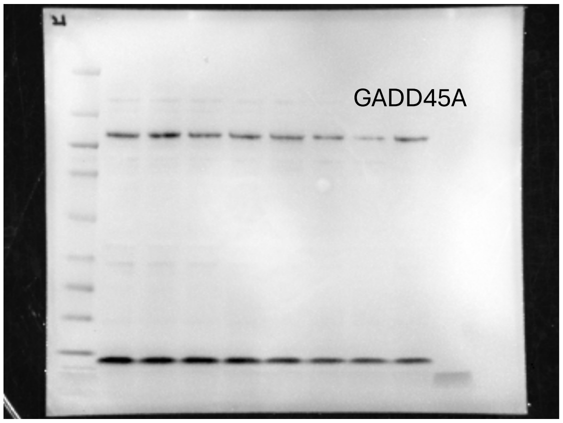


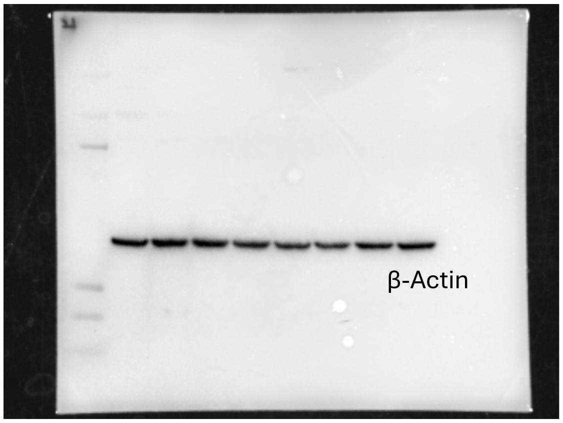


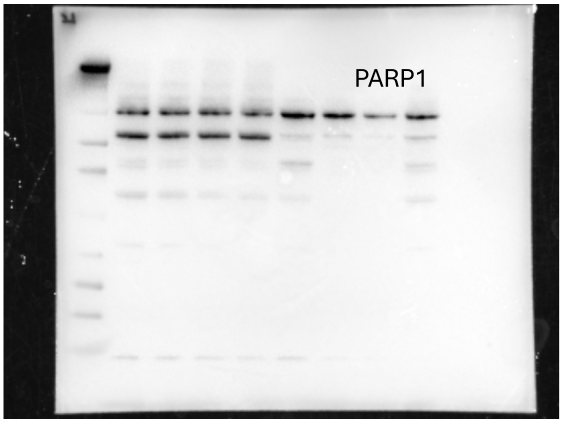


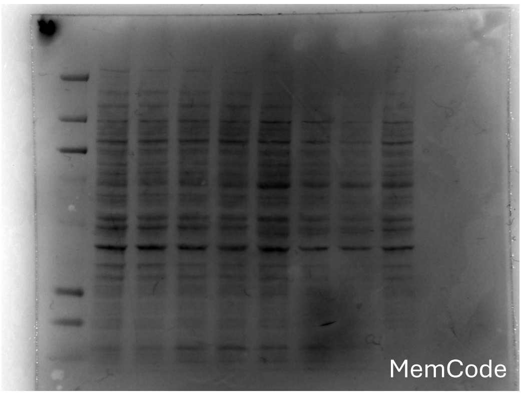

Gel25
Cell: CH22, R-CH22
Workflow: phosphoPERK-Strip/Reprobe-PERK-Strip/Reprobe-β Actin-Memcode

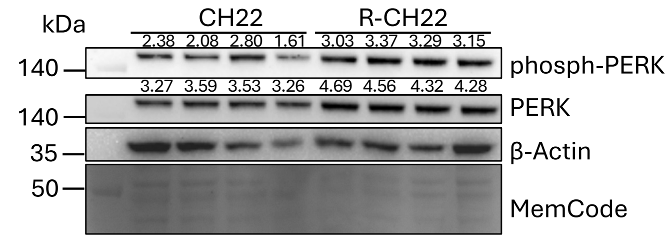


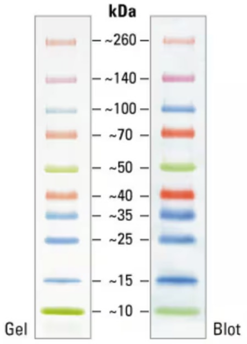


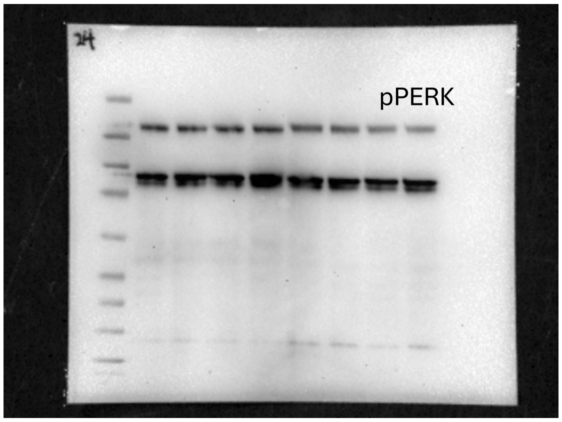


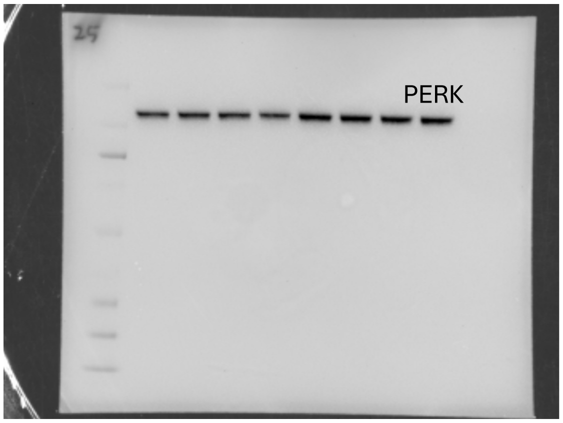


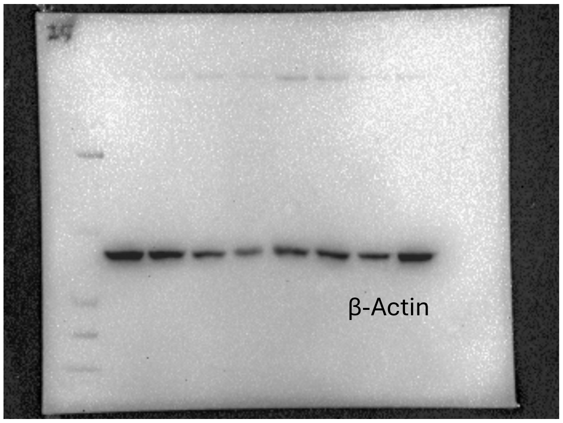


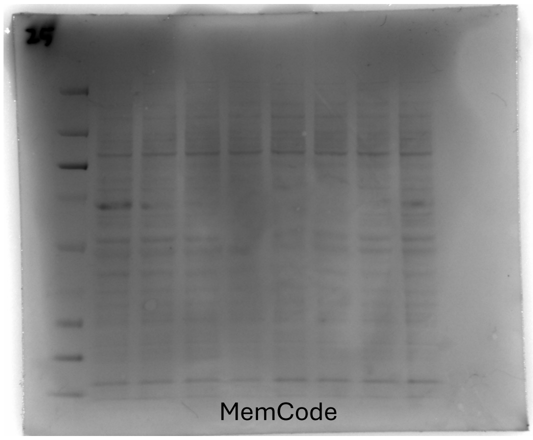

Cell: CH22

Workflow: SREBP1-Strip/Reprobe-FASN-Strip/Reprobe-β Actin-Memcode

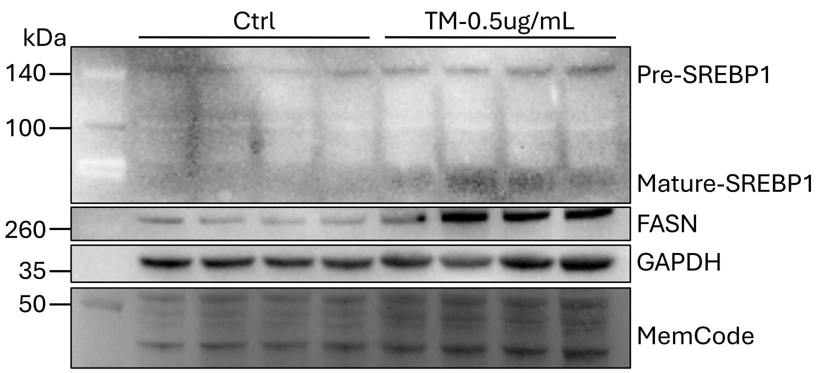


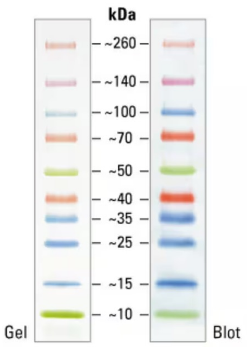


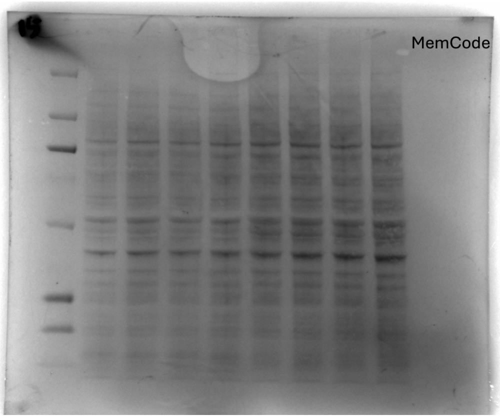


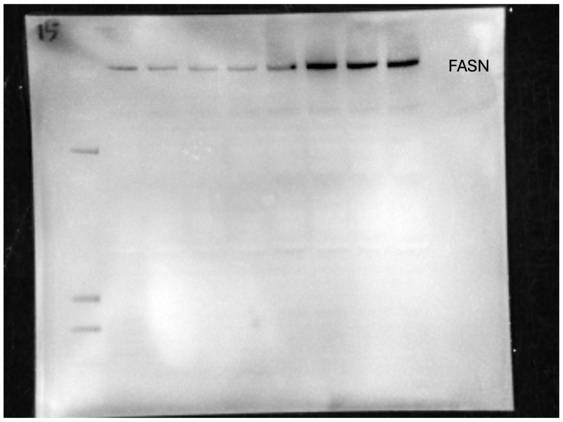


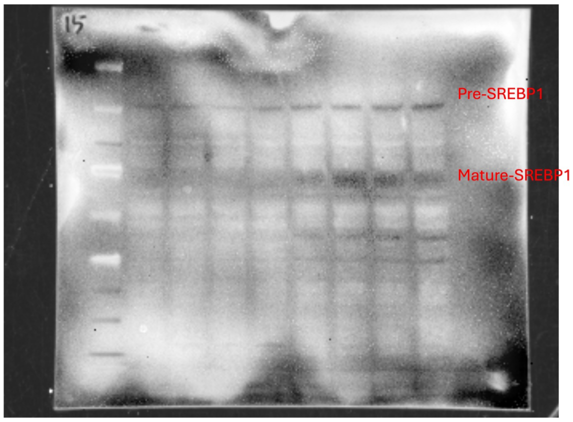


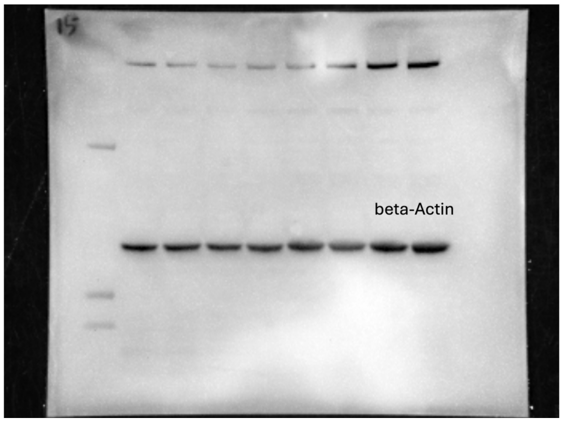

Cell: CH22
Workflow: CSPS3-Strip/Reprobe-PARP1-Strip/Reprobe-β Actin-Memcode

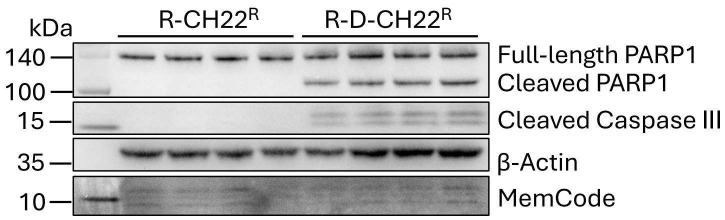


FASNi: TVB-2640, 1.25mM, 48hrs

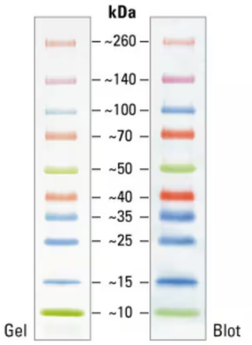


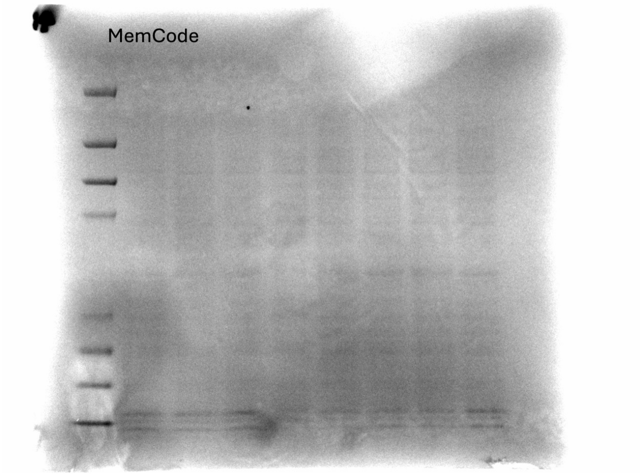


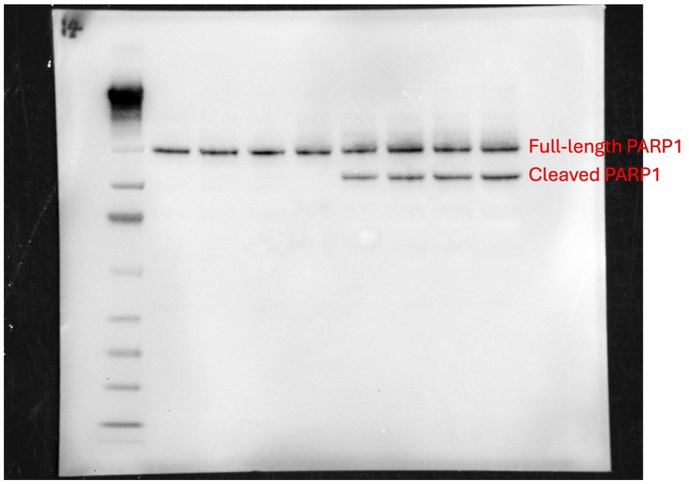


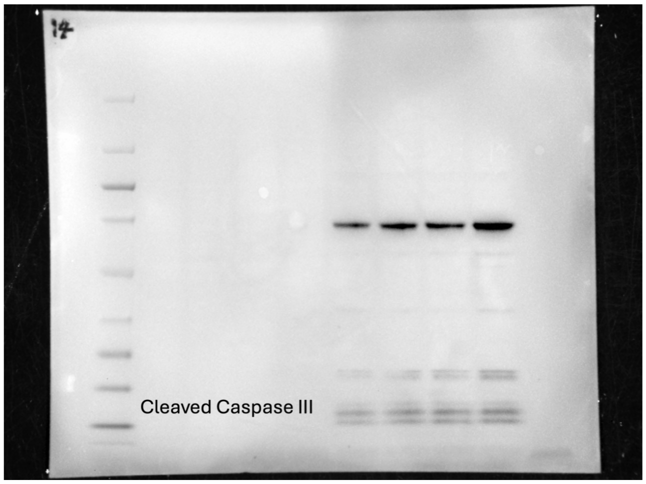


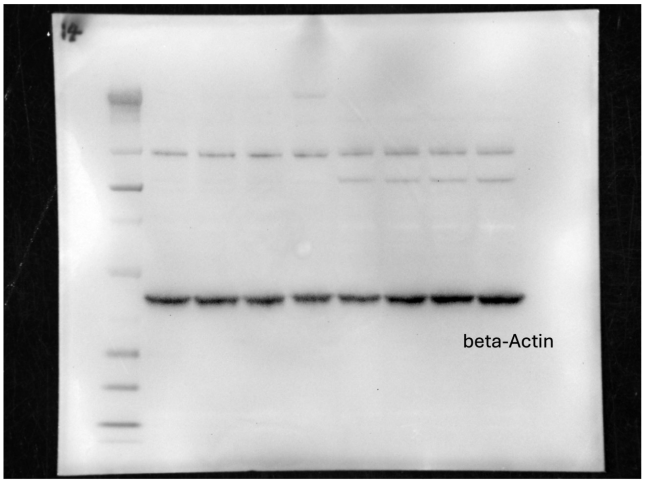

Cell: CH22

Workflow: PARP1-Strip/Reprobe-CSPS3-Strip/Reprobe-β Actin-Memcode

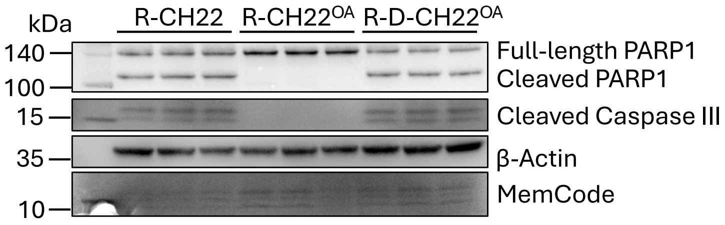


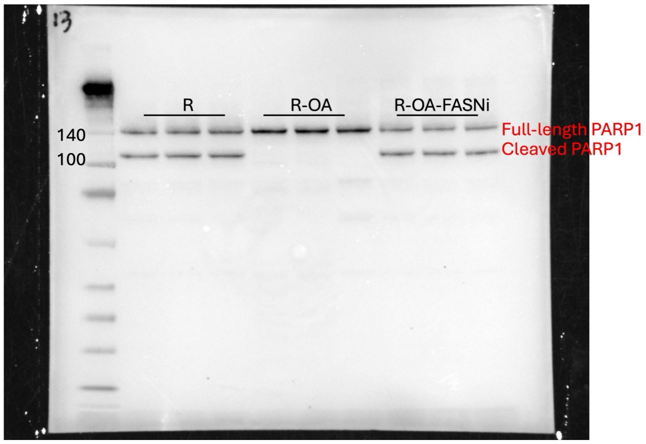


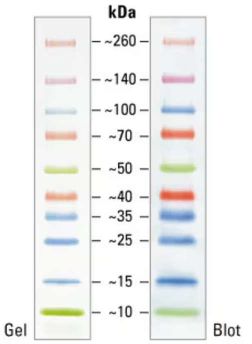


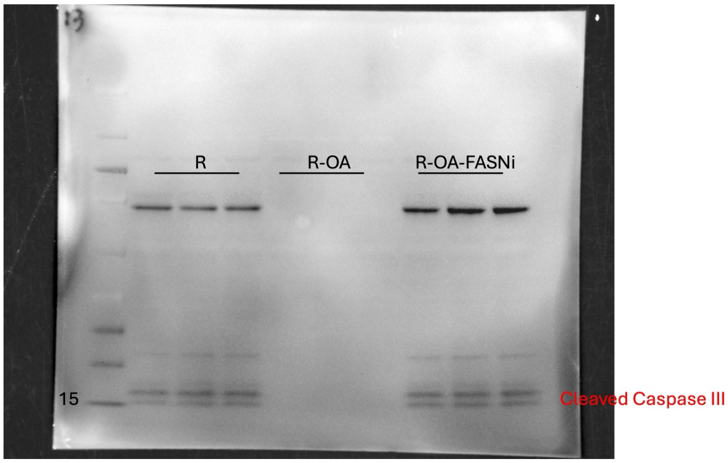


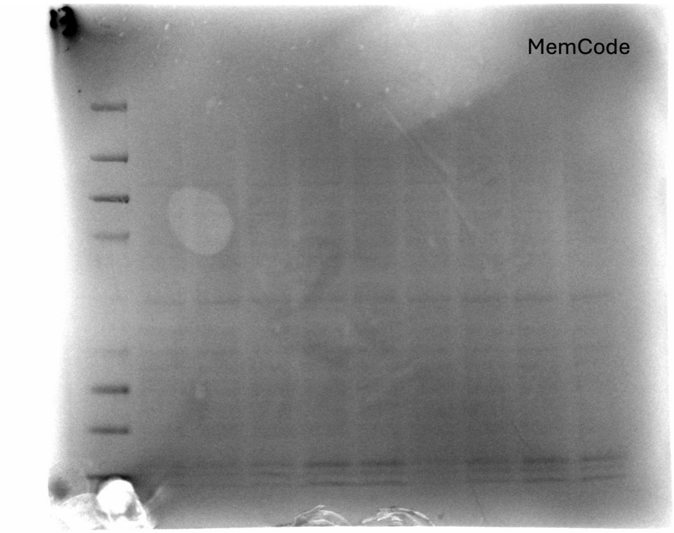


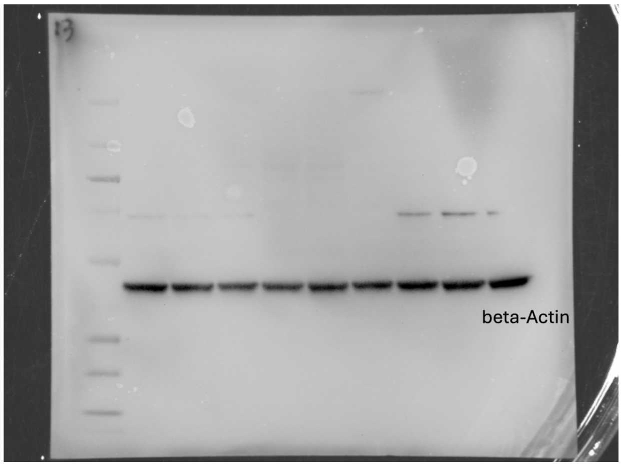
